# Genomic signatures of coevolving phage-bacteria interaction networks

**DOI:** 10.64898/2026.10.08.757702

**Authors:** Marcia F. Marston, Claudia Weihe, Jennifer B. H. Martiny

## Abstract

In diverse natural communities, cells and phages interact with many potential partners, but the impacts of pairwise coevolution on the overall structure of phage-bacteria interaction networks (PBINs) is not fully understood. Here, we conducted multiple pairwise coevolution experiments and examined how coevolution with one partner could simultaneously influence interactions with other partners. We then analyzed the underlying genetic changes that alter interactions in a large cross-experiment PBIN. The results reveal that interactions at the cell surface as well as intracellular interactions can alter patterns across the PBIN. When phages utilized the same set of host cell-surface receptor genes, pairwise coevolution with one partner did alter interactions with other partners. In contrast, if different sets of host cell-surface receptor genes were used, modular patterns in the cross-experiment PBIN were generally observed. On the phage side, resistance-breaking mutations arose in many putative receptor-binding proteins, and these changes could alter interactions with other hosts. However, in both the hosts and phages, over half of the mutated genes were not associated with cell-surface interactions. Host mutations likely altering intracellular phage-host interactions, including translation, protein stability, and DNA replication often led to broad resistance across the PBIN. In phages, mutations occurring in genes encoding endolysins, DNA methylases, oxygenases, transcriptional regulators, and tRNAs were observed potentially also influencing intracellular interactions. Based on these results, we propose a model to illustrate how these two different types of genetic signatures (i.e. extracellular cell-surface interactions verses intracellular interactions) can impact the structure of PBINs.

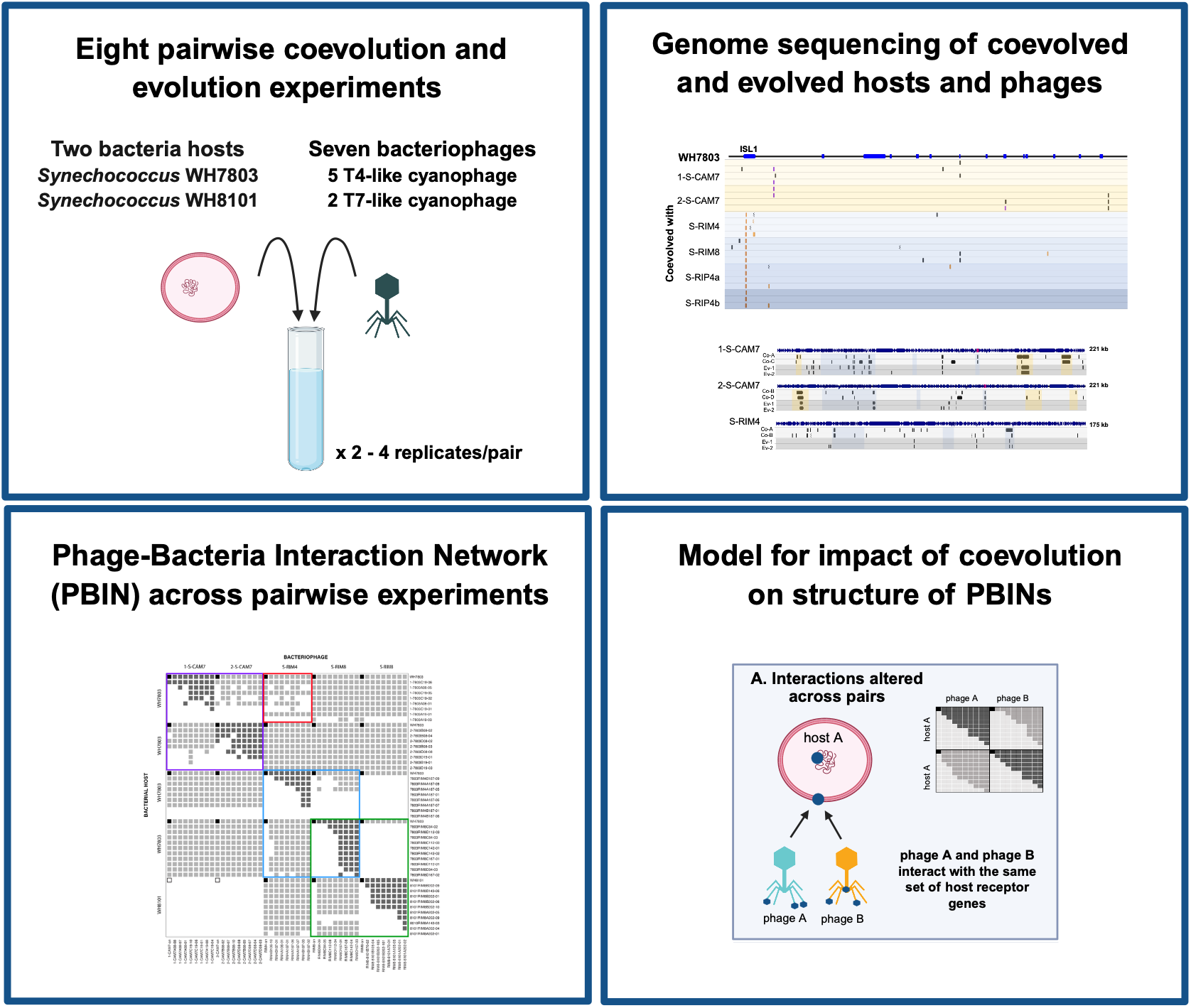
Graphical Abstract.

## Introduction

Unicellular phytoplankton and bacteria drive global biogeochemical cycles in the oceans. These microbial communities are controlled by both “bottom-up” (environmental resources) and “top-down” (predation) factors. An important top-down control of microbial growth are viruses, the most numerically abundant biological entities in the ocean [1]. Unicellular *Synechococcus* and *Prochlorococcus* (Cyanobacteria) are abundant in the oceans and are responsible for up to 25% of ocean net primary productivity [2]. Viruses that infect cyanobacteria (cyanophages) can alter marine carbon and nutrient cycling through the lysis of infected hosts and metabolic reprogramming of cyanobacterial cells during infection [3]. To understand how phages influence bacterial communities, we need to understand phage host ranges (who infects whom), bacterial host sensitivities (who can resist whom), and how these interactions change over time.

Host ranges and phage sensitivities are both extremely variable, leading to a high diversity of interaction phenotypes in a community. For instance, host ranges can be very broad, as in the case of T4-like cyanophages that frequently infect strains across many clades of *Synechococcus* spp. [4], whereas T7-like cyanophages often infect only the *Synechococcus* strain used for isolation [5,6]. While individually documenting every phage-bacteria interaction in the ocean is infeasible, characterizing emergent features of species interaction networks may be tractable (e.g. [7,8]. Indeed, marine phage-bacteria interaction networks are typically nested, where specialist phages tend to infect hosts that are most susceptible to infection [9] (Fig. 1A). Such a pattern may be a signature of coevolutionary dynamics, with directional selection for increased host range and broader resistance [10,11]. In contrast, a large marine infection network was observed to be highly modular [12], such that interactions were more common within particular groups (modules) of phage and bacteria than across groups. In this case, modularity was thought to arise from the geographic scale of the sampled network, where the degree of phage-bacteria interactions was relatively high within sites compared to across sites. Thus, phage-bacteria interaction networks (PBIN) offer a way to characterize and understand the diversity of phage-bacteria interactions in the ocean [13,14].

**Figure 1.**
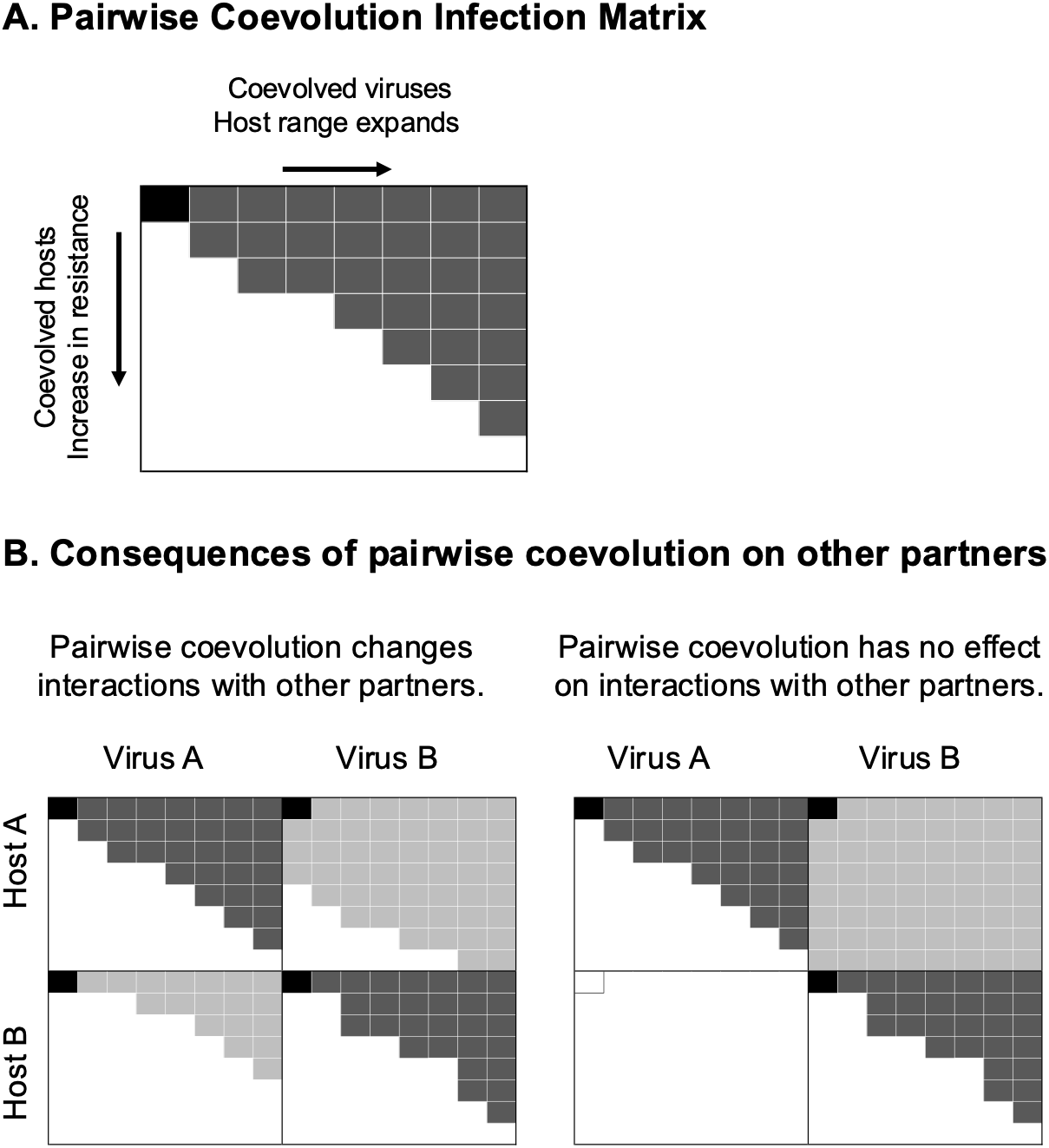
(A) Schematic of a nested pattern of pairwise coevolution. The top left square of the matrix specifies the ancestral interaction, where a black square indicates that the virus can infect the host. Typically, pairwise phage-bacteria coevolution leads to host strains that expand their resistance and phages that expand their host range, such that they can be arranged in a nested, upper-triangular pattern (filled squares indicate infection, white squares indicate resistance). (B) Possible outcomes of coevolution within a phage-bacteria pair on other partners. The left-hand panel illustrates a situation where evolution of increased host resistance (or phage host range) with one partner alters interactions with other partners. Alternatively, coevolution within one phage-host pair may have no effect on interactions with other partners (right-hand panel). This would lead to a more modular interaction network that remains stable despite pairwise coevolution.

The structure of a PBIN will be determined in part by the underlying genetic mechanisms that govern phage-bacteria interactions. However, the difference between whether a phage infects a host can depend on seemingly minor genetic differences. Nearly genetically identical phages can have different host ranges [4], and nearly identical hosts can have differing phage sensitivities [15]. Further, phage-bacteria interaction networks are not static with studies showing that antagonistic coevolution of marine phage and bacteria is ongoing in marine environments [14,16].

Here, we aimed to understand how the structure of coevolving phage-bacteria interaction networks depends on the genetic mechanisms that govern phage-bacteria interactions. Laboratory experiments of interacting phage-bacteria pairs find that antagonistic coevolution leads to minor point mutations and indels that correspond with a nested network structure of the coevolved hosts and viruses [9,17,18]. What is less known, however, is whether such pairwise coevolution generally alters interactions with other potential partners. Certainly, there is evidence of many pleiotropic interactions. The evolution of host resistance to one phage can change the susceptibility of the host to other phages [19,20], and conversely, the evolution of a phage’s ability to infect one bacterial host can alter its ability to infect other hosts [18,21,22]. However, previous studies generally consider interactions from either the host or the phage perspective, rather than how pairwise coevolution simultaneously alters interactions with both other host bacteria and other phages. To do this, we focused on *Synechococcus* and cyanophages as a model system, conducting eight pairwise coevolution experiments with seven cyanophages, including both broad host range T4-like cyanophages and narrow host range T7-like cyanophages and two hosts (*Synechococcus* sp. WH7803 and WH8101). Hosts and phages were isolated throughout each pairwise experiment, and we sequenced the genomes of both hosts and phages to identify mutations that arose during these pairwise interactions. We then conducted infection assays within and across the coevolution experiments to characterize infection patterns.

We addressed three questions: (1) Do the same genes and genomic regions evolve when a host or phage interacts with multiple partners? (2) How does coevolution with one partner alter the interactions with other partners? (3) How do the genomic signatures of phage-bacteria interactions relate to the network structure of these interactions? We hypothesized that the network structure across many phages and hosts would depend on the degree to which virus-host pairs utilize distinct genes and/or genomic regions (Fig. 1B). If the interactions involved a subset of genes shared among most other phages and/or hosts, then pairwise coevolution would likely alter the interactions with many other partners. Alternatively, phage-host interactions might depend primarily on genes that are flexible and are only sporadically present across strains [23,24]. In this case, the potential for a phage and bacterium to interact would depend on the presence of specific genes. We predict that this model would lead to a modular interaction network, where coevolution with one partner only alters interactions with other partners that have the same genes. Of course, many intermediate scenarios might also be possible. In particular, the degree to which the interactions depend on shared genes might differ between phage and bacteria.

## Materials and Methods

### Coevolution and evolution experiments

To identify phage and host interaction genes, we conducted three types of experiments in replicated semi-continuous or continuous culture: (1) pairwise host-phage coevolution, (2) phage evolution on susceptible hosts, and (3) host evolution in the absence of phage (Table 1). In the coevolution experiment, interacting host and phage evolved in response to resistance or infectivity in one another. In the phage evolution experiment, the phage was reintroduced to susceptible hosts at each transfer, allowing the phage to adapt culture conditions and the ancestral susceptible host, but not to cells that had evolved resistance. The host evolution treatment was used to control for adaptation of the host to culture conditions either in a chemostat or flask. Additional details available in supplemental methods. Five T4-like phage isolates (1-S-CAM7, 2-S-CAM7, S-RIM4, S-RIM8, S-RIM44) and two T7-like phage isolates (S-RIP4a and S-RIP4b) obtained from the coastal waters of Rhode Island or California, USA were evolved and coevolved with *Synechococcus* sp. WH7803 and/or *Synechococcus* sp. WH8101 (Table 1, Table S1). The population dynamics and preliminary sequence information for WH7803 coevolved with S-RIM8 have previously been described [25]. As in that initial study, each experiment was initiated with cells derived from a single colony of *Synechococcus* sp. WH7803 or *Synechococcus* sp. WH8101, which were designated as the ancestral host. Likewise, viral isolates were plaque-purified prior to the start of each experiment, and these were designated as the ancestral phage. At selected time points during each of the coevolution and evolution experiments, single-cell colonies and plaque-purified phages were isolated as previously described [19,25,26]. The ancestral, coevolved, and evolved phage and host isolates were subsequently used for genome sequencing and some were selected for host range assays (Datasets S1, S2). In addition to purified isolates, whole communities of cells and/or phages were sequenced in some of the experiments. Across the eight pairwise experiments, we sequenced 80 *Synechococcus* isolates and 12 *Synechococcus* communities (Dataset S1), and 120 phage isolates and 18 phage communities (Dataset S2). See supplementary materials for details on genome sequencing.

**Table 1.** Pairwise coevolution and evolution experiments.

| Virus | Host | Coevolution conditions | Length of coevolution | Evolution conditions | No. of transfers |
| --- | --- | --- | --- | --- | --- |
| 1-S-CAM7_0809CC03 | Syn. WH7803 | Semi-continuous | 19 transfers | Semi-continuous | 19 |
| 2-S-CAM7_0910SB42 | Syn. WH7803 | Semi-continuous | 19 transfers | Semi-continuous | 19 |
| S-RIM4_RW_11_0999 | Syn. WH7803 | Chemostat | 187 days | Semi-continuous | 25 |
| S-RIM8_RW_22_0300 | Syn. WH7803 | Chemostat | 167 days | Semi-continuous | 25 |
| S-RIM8_RW_22_0300 | Syn. WH8101 | Chemostat | 202 days | Semi-continuous | 25 |
| S-RIM44_Np_20_0711 | Syn. WH8101 | Chemostat | 202 days | Semi-continuous | 25 |
| S-RIP4a_RW_07_0909 | Syn. WH7803 | Chemostat | 170 days | Semi-continuous | 24 |
| S-RIP4b_RW_20_0813 | Syn. WH7803 | Chemostat | 170 days | Semi-continuous | 24 |

<sup>a</sup> Semi-continuous culture conditions consisted of flasks with 20 mL seawater medium, cells, and phage. For coevolution, an aliquot of the previous week’s culture was transferred into a new flask with fresh media. For evolution experiments, phage was reintroduced to susceptible hosts at each transfer.
<sup>b</sup> Chemostats provided continuous culture conditions where seawater medium was supplied to each 35-mL chemostat containing cells and phage at a dilution of ~1.02/d. (See details in Supplementary Materials).

### Genomic analyses

The sequences from the ancestral cell and phage isolates used to initiate each co/evolution experiment were used as references to identify genetic changes that occurred during the experiments (Table S1). Illumina MiSeq reads were trimmed, mapped back to the ancestral genome, and SNPS and indels were identified. (Details in supplemental methods). For host cell and phage isolates, only SNPs or indels with >80% of reads supporting the mutation were included in the final analyses. For cell and host communities, all SNPs/indels with >10% supporting reads were considered. (See Datasets S3 through S10 for all host and phage mutations identified).

The Cyanorak database [27] was used to determine if the observed mutations in *Synechococcus* hosts occurred in genes that were core (present in all fully sequenced *Synechococcus* in the database) or flexible (only present in some *Synechococcus*) and then to examine if these genes were located in previously defined genomic islands. The predicted functions of *Synechococcus* genes were identified using the Cyanorak database of manually curated ‘clusters of likely orthologous genes’ (CLOGs). DefenseFinder was used to scan for known phage defense systems in *Synechococcus* strains WH7803 and WH8101 and for anti-defense systems in the phages used in this study [28–30]. In the phages, the assignment of genes as core or flexible was difficult due to the divergence of the two CAM7 strains from the other T4-like cyanophage (<10% genome-wide ANI; Fig. S1) [31]. Thus, to facilitate the comparison of mutated genes among the phage isolates, we reannotated the phage genomes using Pharokka [32], which assigned predicted proteins into PHROG groups (clusters of viral protein families generated by using a remote homology detection by HMM profile-profile comparisons [33]). This enabled us to compare mutated phage genes across the CAM and RIM strains based on PHROG functional annotations (e.g. tail-associated proteins), rather than relying on sequence homology.

### Infectivity assays and network analyses

We first conducted infectivity assays within an experiment (one host-phage pair). Ancestral and coevolved *Synechococcus* isolates were challenged with ancestral and coevolved phages. All assays were carried out in duplicate in 24– or 48-well microtiter plates as previously described [19,25]. Control wells containing *Synechococcus* cells, but no phage, were included on each plate. Wells were monitored visually for cell lysis for up to four weeks. *Synechococcus* cultures were scored as susceptible (and the virus as infective) if growth was visibly inhibited compared with the control cells-only wells. Cells that were not lysed after 4 weeks were considered to be resistant to the phage [25].

Five coevolution experiments resulted in a diversity of phage infectivity and host sensitivity phenotypes. From each of these five pairwise experiments, we then selected at least four hosts and four phages from two replicates to include in cross-infection assays across experiments. The isolates were chosen to represent a diversity of resistance and infectivity patterns observed within each experiment (i.e. we selected isolates representing different infection/sensitivity phenotypes). Isolates from three additional pairwise experiments were not used for this cross-infection assay. In one case, the coevolved phage did not produce distinct plaques for isolation (for the WH8101-S-RIM44 pair), and in two other cases, the coevolved T7-like phages did not evolve to overcome host resistance, resulting in limited phenotypic diversity in the WH7803-S-RIP4a and WH7803-S-RIP4b pairs. Notably, the lack of resistant-breaking mutations in coevolved T7-like cyanophages was also observed in a previous study [34].

The final infection matrix consisted of 49 cells and 45 phages across five pairwise coevolution experiments. We first tested whether interactions between isolates from each pairwise coevolution experiment were significantly nested using the NODF metric, which ranges from 0 (not nested) to 1 (perfectly nested) and a Bernoulli null model (setting = Equiprobable) with 10,000 randomizations using the BiMat package for MATLAB [9].

In some cases, pairwise coevolution did not change any interactions with partners from another experiment – that is, all interactions were the same as the ancestor (as in Fig. 1B). Such cases were clearly modular. In the cases where pairwise coevolution resulted in changes in interactions with other partners (ancestral and/or coevolved strains), we then tested whether interactions with new partners were nested using the same procedure described above. Note that because we wanted to test for modularity of particular modules based on *a priori* knowledge, we did not aim to classify the interactions into *de novo* modules (as in [9]).

## Results and Discussion

### Genomic signatures of *Synechococcus-*phage interactions

#### Interaction genes in *Synechococcus*

We sequenced 56 *Synechococcus* WH7803 isolates that had coevolved with one of 6 phages and 15 *Synechococcus* WH8101 isolates that had coevolved with one of two phages. Overall, in the coevolved hosts, 34 protein-encoding genes and six intergenic regions (IGR) were mutated in WH7803 (Table S2) and 24 protein-encoding genes and one IGR in WH8101 (Table S3). Most changes in protein-encoding genes (88% in WH7803; 94% in WH8101) were nonsynonymous and led to amino acid substitutions, frameshifts, premature stop codons, or removal of stop codons. These changes in the coevolved hosts are likely due to interactions with the phage, rather than adaptation to the laboratory conditions, as there was no overlap between these mutations and the few mutations observed in evolved (no-phage) WH7803 or WH8101 controls (Datasets 3 and 4).

Cell-surface interactions between bacteria and phage are important determinants of host range and mutations that block phage absorption can lead to resistance (reviewed in [35,36]). In this study, we observe that 85% of coevolved WH7803 and WH8101 host isolates (60 out of 71) acquired at least one, and often several, mutations in genes involved in the synthesis or modification of cell-surface receptors (Table S4, Datasets 3 and 4). Across all coevolved *Synechococcus* isolates, mutations occurred in 18 different cell surface receptor-associated genes in WH7803 and 5 such genes in WH8101 (Table S4). Only 3 of these 23 receptor-associated genes (i.e., *galE, rfbA,* and *wcaJ*) contained mutations across phage-host pairs and none of the same genes mutated in both WH7803 and WH8101, even though almost all mutated surface-receptor genes have homologs in both host isolates (Tables S2, S3).

In natural communities, single-cell sequencing of *Prochlorococcus* spp. revealed that nearly identical strains often possess many differences in outer membrane modification genes located in genomic islands [37]. The high turnover of genes in genomic islands has been attributed to selection pressure to escape phage infection [20,23]. Previous laboratory coevolution studies of cyanophages with *Synechococcus* or *Prochlorococcus* hosts identified resistant-conferring mutations in cell-surface receptors genes located in genomic islands [23,25]. Likewise in this study, many of the receptor-associated genes containing mutations in coevolved WH7803 isolates were in a single genomic island (ISL1) (Fig. 2). In an earlier study examining WH7803-S-RIM8 pairwise coevolution, we identified mutations in two surface-receptor related genes (*rfbA* and a glycotransferase-encoding gene) in genomic island 1 (ISL1) [25]. Here, genomic sequencing of additional coevolved WH7803 isolates from that study, as well as WH7803 isolates coevolved with 1-S-CAM7, 2-S-CAM7, S-RIM4, S-RIP4a and S-RIP4b, revealed changes in 11 genes and 2 intergenic regions in ISL1. Notably, a single mutation in the upstream start region of the *rfbA* gene was observed in 19 WH7803 isolates coevolved with S-RIM4, S-RIM8, or S-RIP4 and conferred resistance to both T4-like myoviruses and T7-like podoviral isolates (Fig. 2, Table S4). The *rfbA* gene encodes an enzyme (glucose-1-phosphate thymidylyltransferase) important in the synthesis of the O antigen of LPS, a predicted receptor for cyanophage attachment [38]. Changes in *rfb* genes have been also shown to affect adsorption in other phage-bacteria systems [e.g. [39]]. While this single genomic island in WH7803 contained over a dozen mutated genes that alter phage-host interactions, it was the only genomic island identified in WH7803 or WH8101 that contained multiple mutated genes (Fig.2). Overall, the majority of mutated genes, including some that are predicted to be involved in synthesis of cell-surface receptors, were located outside of genomic islands (65% in WH7803 and 84% in WH8101) (Tables S2, S3).

**Figure 2.**
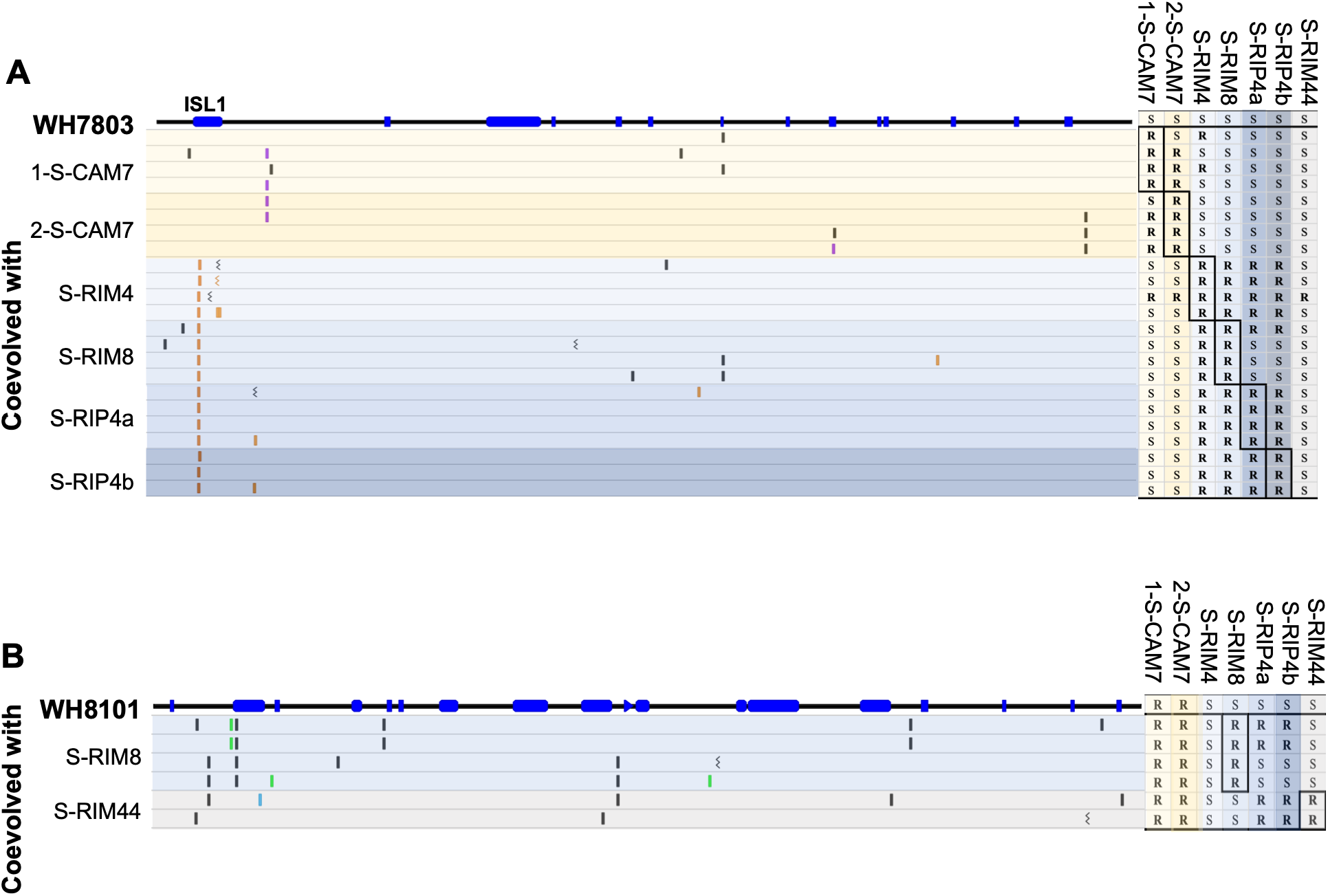
Mutations in coevolved *Synechococcus* hosts. The top line in each diagram represents the host genomes (A) WH7803 (2.36 mb) and (B) WH8101 (2.62 mb). The blue rectangles along the genome are previously described genomic islands [27]. Each row under the host genome is a single cell isolate that was coevolved with the phages listed to the left. The dashes represent single polymorphisms relative to the ancestral sequence. The color of the dashes indicates whether the mutation was in a cell-surface receptor-associated gene (colored) or in a gene not known to be associated with cell-surface interactions (black). Different colors represent mutually exclusive sets of receptor-related genes. Genomic island 1 in WH7803 (labeled ISL1 in A) was the only genomic island that contained mutations in multiple genes across phage-host pairs. The grid on the right shows the host range results for the ancestral host and each coevolved host against the ancestral phages used in the experiments. Resistance to one phage strain often led to resistance to other phage strains and in some cases to all ancestral phage strains in the assay. S, susceptible; R, resistant.

In WH7803 isolates coevolved with the two narrow host range T7-like podoviruses (S-RIP4a and S-RIP4b), mutations were only observed in genes associated with the synthesis of cell-surface receptors. In contrast, over half of mutated genes in hosts coevolved with broad-host range T4-like viruses were not related to cell-surface receptors and likely instead contribute to intracellular mechanisms of phage defense (Table S4). The number of known intracellular defense mechanisms in bacteria is expanding rapidly (reviewed in [40]) and recent studies suggest that up to 1.5% of bacterial genes may be involved in phage defense with many single-gene defenses yet to be discovered [28,41]. *Synechococcus* WH7803 and WH8101 do not possess CRISPR-Cas systems, however, marine *Synechococcus* spp. have been shown to possess intracellular resistance mechanisms that can block multiple steps of phage infection within the host cell [42]. The genetic and molecular mechanisms of these defense systems in marine cyanophages are largely uncharacterized (but see, [43]). None of the mutated *Synechococcus* genes in this study was predicted to be part of a known phage defense system (based on DefenseFinder version 2.0.0 [29]), nevertheless the functional annotations for many of these genes (e.g. proteases, tRNA synthetases, DNA ligase and primase; Table S4) suggest that they could play a role in intracellular defense against phage infection [41,44,45]. For example, across all the pairs in this study, mutations were found in three core *Synechococcus* genes involved in ribosome biogenesis (two in WH7803 and one in WH8101), with mutations in one of these genes conferring resistance to almost all (40 of 45) phages in the PBIN (Table S4). Additionally, mutations in three different core tRNA synthetase genes (one in WH7803 and two in WH8101) arose, each in a different viral-host pairs (Table S4). The modification and/or expression of host tRNAs have been shown to influence phage-host interactions in *Synechococcus* [43] and in *E. coli* [45]. Finally, mutations in a DNA ligase gene in WH7803 occurred in several phage-host pairs and a single mutation in this gene could confer resistant to the S-CAM7 ancestral phage. Further molecular characterization is needed to understand the mechanisms of host resistance provided by these mutations.

#### Interaction genes in cyanophages

Next, we examined the genetic changes that occurred in six cyanophage strains when they interacted with *Synechococcus* WH7803 or WH8101. In all, 93 coevolved phage isolates (63 T4-like myoviruses and 30 T7-like podoviruses) were sequenced across seven phage-host pairs. From the phage evolution experiments (where phages were reintroduced to susceptible ancestral hosts each transfer), we sequenced 20 phage isolates (16 T4-like and 4 T7-like) and 18 community lysates (Dataset S2). Within a pairwise coevolution experiment, phages collectively possessed mutations in 13 to 20 regions (regions here refer to single genes, intergenic regions or blocks of adjacent deleted genes). We also observed large numbers of mutated regions in the evolved phages (5 to 20 mutated regions), but the number of mutations in coevolved phages always outnumbered those observed in the corresponding phage evolution experiments (Fig. 3, Datasets 5 to 10). As with the host mutations, most of the observed phage mutations were nonsynonymous (73-96% in coevolved pairs and 85–95% in evolved pairs), suggesting positive selection in both the coevolution and evolution experiments as observed in other phage-bacterial systems [21,22,46]. As in these previous studies, we observed overlap in mutated regions among evolved and coevolved phage isolates of the same host-phage pair, particularly in tail-associated proteins (Tables S5 to S9). Thus, adaptation to the *Synechococcus* hosts is likely occurring in both coevolution and evolution experiments which prevented us from easily distinguishing between mutations involved in initial adaptation to the host and those responsible for overcoming host resistance (Fig. 3).

**Figure 3.**
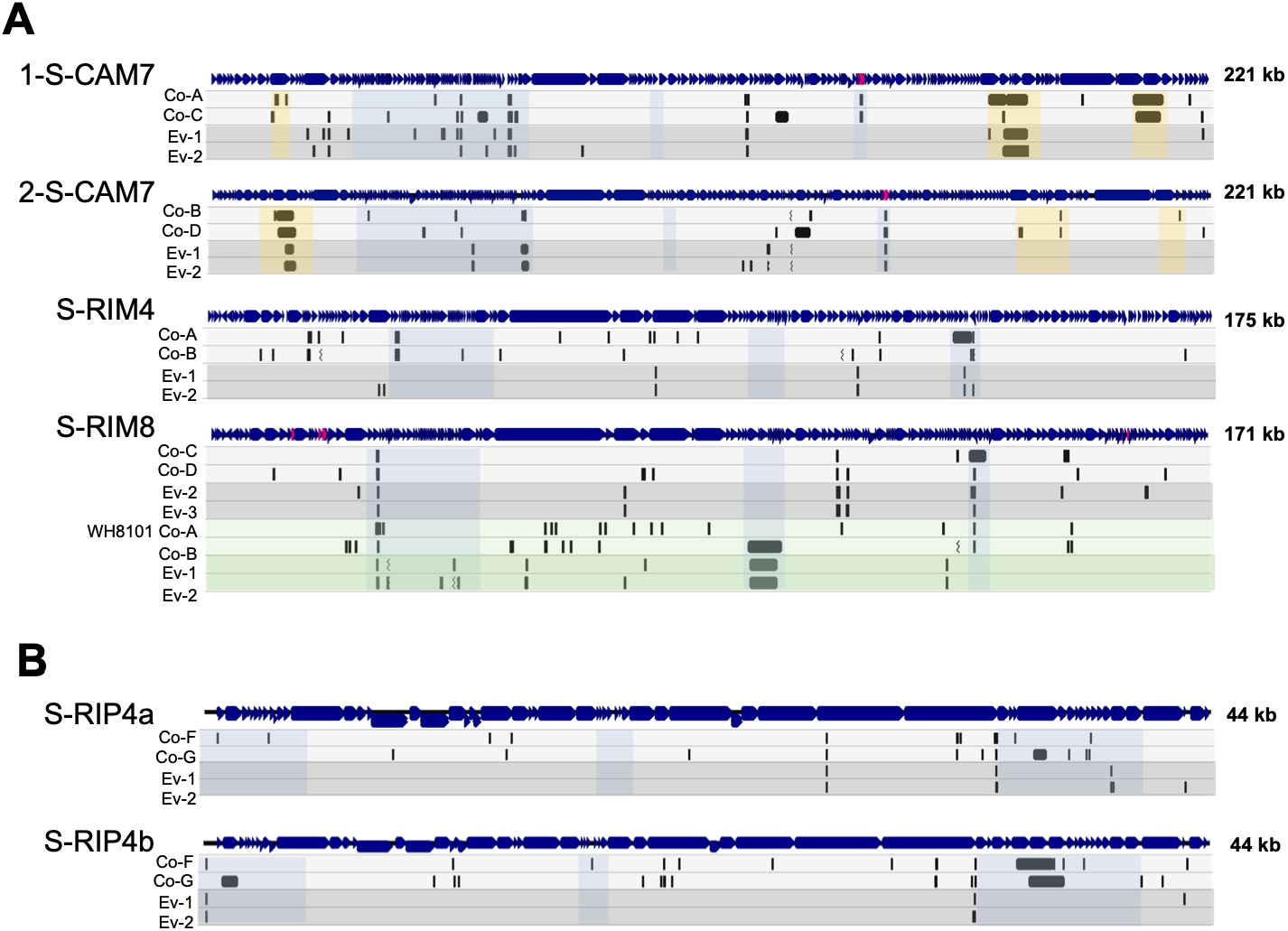
Summary of genetic changes in coevolved and evolved T4-like phage (A) and T7-like phage (B). The genome map of the ancestral phage is shown with ORFs (dark blue) and tRNAs (red). Every phage strain was coevolved (light grey shaded row) and evolved (dark grey shaded row) with *Synechococcus* WH7803. S-RIM8 was also coevolved and evolved with *Synechococcus* WH8101 (light and dark green shaded rows respectively). Dashes represent SNPs and black rectangles indicate large deletions relative to the ancestral genome. The combined mutations found in all sequenced isolates from two coevolved replicates (labeled Co-replicate letter) and two evolved communities (Ev-replicate number) are shown under the ancestral genome map. Community lysates were used for the evolved experiments due to the low number of evolved isolates individually sequenced. Yellow shading in the S-CAM7 phage represents regions that are variable within the S-CAM7 strain and are not present in the other phages. Blue shading indicates variable genomic islands previously identified in T4-like cyanophages (A) [24,53,54] and T7-like cyanophages (B) [55,56].

There were no mutated homologous genes in common among the T4-like phage strains across the pairwise experiments, in part due to the divergence among S-CAM and S-RIM strains (Fig. S1). However, when functional annotations are considered, many of the mutated genes in the T4-like phages (40–56% in coevolved pairs and 36-60% in evolved pairs) are predicted to encode structural proteins (e.g. baseplate, capsid, tail and tail fiber proteins) (Tables S5 to S7; Datasets 5 to 8). In the T7-like phages, over 80% of mutations were in genes encoding structural proteins (Tables S8 and S9; Datasets 9 and 10). Among closely related phages (e.g. the two S-CAM7 strains and the two S-RIP4 strains), there were overlaps in mutated tail fiber genes, but in general, different combinations of phage genes and/or locations within a gene were mutated in each experiment and even among replicates of the same phage-host pair (Fig. 3, Datasets 5 to 10). Similar results were observed in the experimental evolution of T4 bacteriophage on *E. coli* K12 and C host strains, where most mutations occurred in T4 structural genes, but the specific mutations were often unique to a particular population or replicate [21].

Beyond structural genes, mutations were observed in phage genes utilized during infection processes within a cell. Although no known anti-defense systems were detected in these cyanophages using DefenseFinder, some of mutated phage genes may encode proteins to counteract defenses in the hosts (reviewed in [47,48]). Alternatively, these mutations may be due to adaptations to the host. Coevolved phage isolates of 1-S-CAM7 and 2-S-CAM7 possessed mutations in genes encoding a DNA methylase, peptidase, and transcriptional regulator as well as the loss of 3 to 4 adjacent tRNA genes (Table S5). S-RIM4 and S-RIM8 coevolved isolates had mutations in genes encoding endolysins and 2OG-Fe(II) oxygenases (Tables S6 and S7). In coevolved S-RIP4a and S-RIP4b isolates, mutations were observed in the *nblA* gene, involved in the degradation of the host’s phycobilisomes [49] and a ribonuclease reductase gene (Tables S8 and S9). Finally, mutations occurred in many viral genes with unknown functions. Large deletions also were observed in both evolved and coevolved phage isolates and were often in variable regions of the genomes (Fig. 3). More than half of the genetic changes in 1-S-CAM7 and 2-S-CAM7 evolved and coevolved isolates were deletions and many of these were in regions of their genomes that are not shared by the other phage strains (Fig. 3). Additional experimental evidence is needed to determine the effects of these changes and large deletions on phage infection and fitness.

### The effect of coevolution on the phage-host interaction network

Infection assays confirmed that within pairwise coevolution experiments, host resistance and phage infectivity generally evolved in a directional manner [11]. Resistance of the host isolates and host range of the phage isolates generally increased, despite the small number of isolates tested from each chemostat experiment. This dynamic resulted in significantly nested patterns of infectivity within experiments (dark gray squares along the diagonal in Fig. 4; NODF = 0.63-0.77, p<0.001). The one exception was the 2-S-CAM7 and WH7803 experiment, but even in this case, the infection pattern was marginally nested (NODF = 0.66, p=0.08).

**Figure 4.**
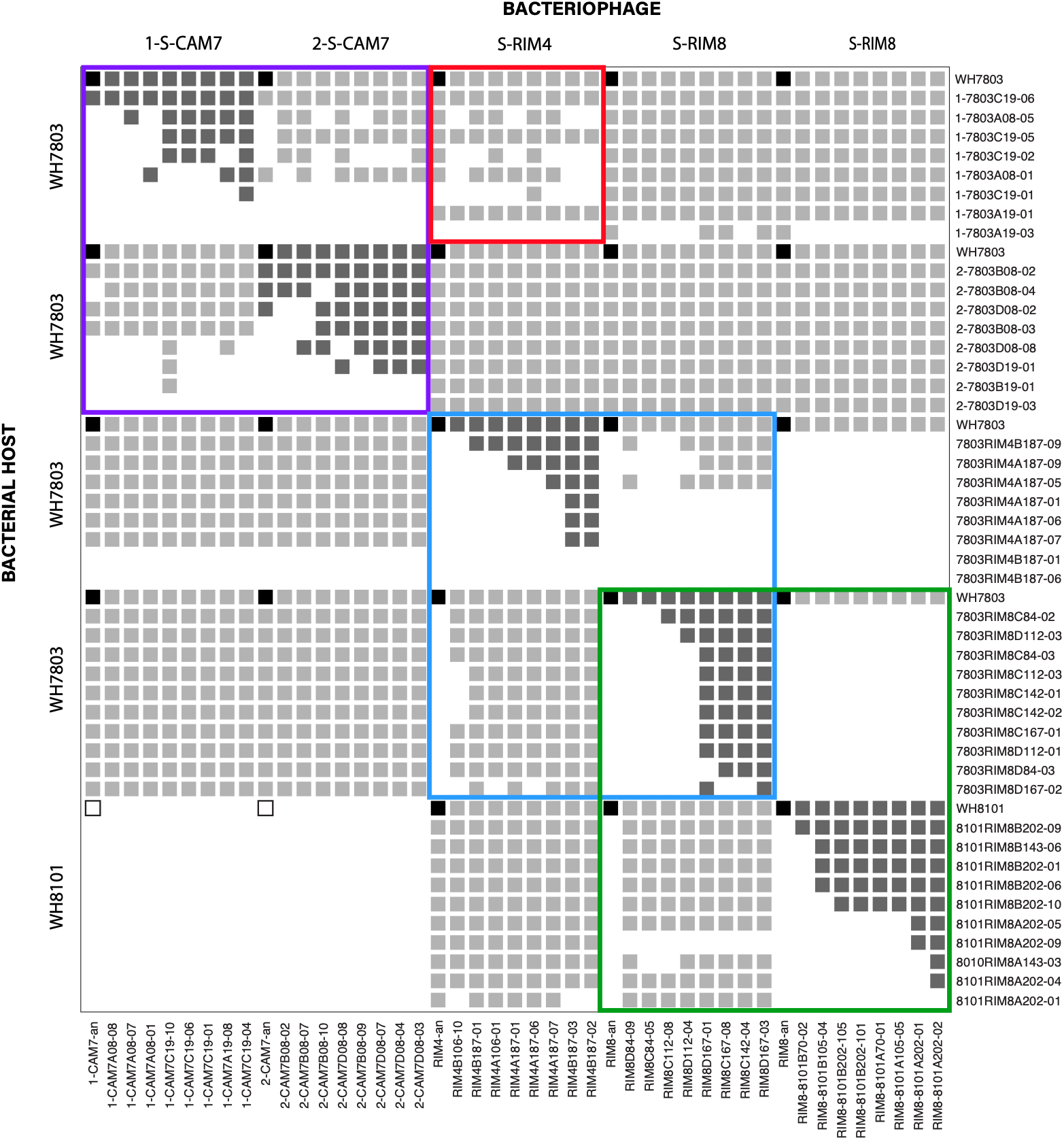
Phage-bacteria interaction network based on five pairwise coevolution experiments (blocks of dark grey on the main diagonal). The ancestral interaction between each host and phage is represented in the top lefthand corner of each of the 25 modules, where a black square indicates that the ancestral phage can infect the host. Between eight and ten coevolved hosts and eight coevolved phages were isolated from each experiment, and all interactions between isolates within and across experiments were recorded. As in Fig. 1, filled squares indicate infection, and white squares indicate resistance. See text for explanation of colored outlines.

We next investigated if pairwise coevolution altered interactions with non-coevolving partners by conducting infection assays across experiments. In contrast to studies that aim to identify interaction modules [12], here we defined a module *a priori* based on the ancestral strains. Thus, Figure 4 illustrates 25 modules, or blocks of potential interactions. In the first four of the five experiments, the ancestral host (WH7803) was the same, allowing us to compare the effects of coevolution with different virus partners.

Generally, coevolution with one partner (blocks of dark grey on the main diagonal) greatly altered interactions with one or two other modules (those off the main diagonal) (Fig. 4). For instance, consider the 1-S-CAM7 and 2-S-CAM7 coevolution experiments (outlined in purple in Fig. 4). Coevolution of 1-S-CAM7 with WH7803 led to six different 1-S-CAM7 host range phenotypes, based on infection patterns within the coevolution experiment. In turn, these phenotypes varied in their ability to infect WH7803 isolates that had coevolved with 2-S-CAM7. The reciprocal pattern was also true; 2-S-CAM7 isolates varied in their ability to infect WH7803 isolates that coevolved with 1-S-CAM7. A similar, reciprocal pattern was also observed for RIM4 and RIM8 when they were coevolved with WH7803 (outlined in blue in Fig. 4).

In the fifth experiment, WH8101 was used as the host, allowing us to consider the effects of coevolution with the same phage (S-RIM8) on two different hosts (outlined in green in Fig. 4). In contrast to the experiments that held the host constant, here the infectivity patterns were not reciprocal. WH7803 isolates that had coevolved with S-RIM8 were resistant to all the S-RIM8 isolates that have coevolved with WH8101. In contrast, WH8101 isolates that coevolved with S-RIM8 remained largely sensitive to S-RIM8 isolates coevolved with WH7803. Thus, not only did coevolution alter interactions with other potential partners, but in this case, the outcome of the interactions was dependent on which host the phages coevolved with first. A non-reciprocal pattern was also observed between the S-RIM4-WH7803 versus S-RIM8-WH8101 experiments.

Yet beyond pairs discussed above, coevolution did not alter interactions with most partners, leading to modular blocks of light gray or white in Fig. 4. For example, coevolved isolates of 1-S-CAM7 and 1-S-CAM7 infected nearly all WH7803 isolates that coevolved with S-RIM4 and S-RIM8. Conversely, WH7803 and WH8101 isolates that coevolved with S-RIM8 did not change their interactions with 1-S-CAM7 or 2-S-CAM7. Two notable exceptions occurred, however. First, many WH7803 isolates coevolved with 1-S-CAM7 did alter their interactions with RIM4 (outlined in red in Fig. 4). Second, two WH7803 isolates that coevolved with S-RIM4, became resistant to all phages (i.e., the two white rows across the middle of the matrix).

### Connecting genomic signatures with the interaction network

The interactions between bacterial cell-surface receptors and phage receptor binding proteins are a key determinant of infection, and recent studies have shown it is possible to predict infection outcomes based on clade specific bacterial-surface receptors and phage receptor-binding proteins [14,50]. In this study, when the phage strains interacted with the same host gene (or set of genes) encoding cell-surface receptors, coevolution within a phage-bacteria pair altered interactions with other partners (Fig. 5A). For example, in the two S-CAM7 strains (outlined in purple in Fig. 4), mutations in a UDP-glucose 4-epimerase (*galE*) gene (involved in LPS production) occurred in WH7803 cells in all replicates of both sets of pairwise experiments, although the exact location and types of mutation varied across replicates (Dataset S3). Mutations in this gene provided resistance to both ancestral S-CAM7 strains as well as some coevolved S-CAM7 isolates. Resistance-breaking mutations did occur in S-CAM7, likely due to mutations in structural or tail fiber genes that allow attachment and infection to occur (Tables S5).

**Figure 5.**
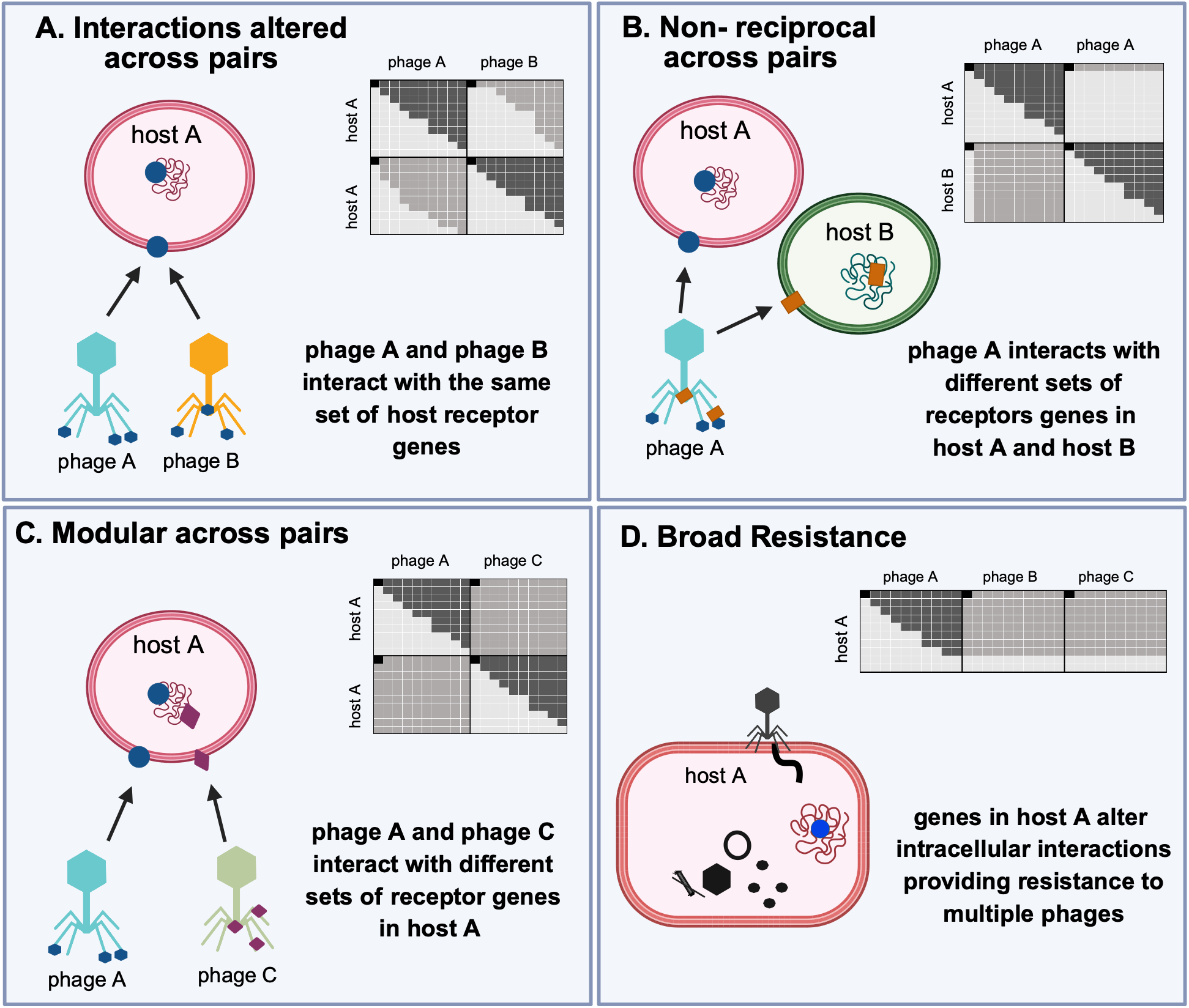
Genomic signatures of phage-bacteria interaction networks. In each diagram, shapes of the same color represent sets of genes in the hosts that change specific cell-surface receptors and the corresponding phage structural proteins interacting with these receptors. In the infection matrixes, blocks of dark grey on the diagonal are the results of pairwise coevolution experiments. Blocks with light grey or white illustrate the consequences of pairwise coevolution on other partners. As in Fig. 4, filled squares indicate infection, and light squares indicate resistance. **A.** Coevolution between phage-bacteria pairs altered interactions between other partners when phage strains interacted with the same host gene (or set of genes) encoding cell-surface receptors. **B.** Non-reciprocal interactions were observed when a phage used some of the same virion structural and tail fiber genes to interact with different sets of receptors on Host A and Host B. In this case, phage A isolates coevolved on host A can infect all coevolved host B isolates, but not vice versa. **C.** Modular interactions were observed when phage interacted with different cell surface receptor genes and thus coevolution with one partner had no effect on interactions with other partners. **D**. Some coevolved host isolates possessed mutations in genes that led to broad resistant to many different phage strains. This is likely due to intracellular resistance mechanisms that prevent the production of new phage. Figure created in https://BioRender.com

Likewise in pairwise coevolution between WH7803 and S-RIM4 or S-RIM8 (outlined in blue, Fig. 4), all host isolates resistant to the ancestral phages possessed at least one mutation in a single operon in a genomic island encoding genes in the LPS biosynthesis pathway (Fig. 2; Table S2). Some S-RIM4 and S-RIM8 isolates evolved to overcome this resistance, but due to the large number of mutations in coevolved phages, it is not known if the resistance-breaking mutations were in the structural genes shared among S-RIM4 and S-RIM8 isolates, or flexible genes that only appear in S-RIM4 or S-RIM8. Interestingly, here the interaction is asymmetrical, with very few coevolved S-RIM8 isolates able to overcome the resistance in WH7803 cells coevolved with S-RIM4. One possible explanation is that most resistant WH7803 isolates coevolved with S-RIM4 contain mutations in multiple genes within ISL1, whereas the WH7803 isolates coevolved with S-RIM8 have mutations in only a single gene in ISL1 (Table S2; Datasets S7 and S8). Mutations in different parts of the cell-receptor biosynthesis pathway may influence the likelihood of resistant-breaking phage isolates arising. Further characterization is needed to assess whether S-RIM8 isolates can develop resistance-breaking mutations to infect WH7803 isolates coevolved with S-RIM4 or if S-RIM8 phages have structural limitations.

In a few cases, non-reciprocal interactions were observed, and these were difficult to predict. When WH7803 and WH8101 were coevolved with the same phage (S-RIM8) (outlined in green in Fig. 4), resistance was linked to different sets of surface-receptor related genes in each host (Fig. 5B). In WH7803, resistance to S-RIM8 phage was associated with mutations in *rfbA* and *rfbD* genes, whereas in WH8101, resistance was correlated with mutations in *ksdD* and *lptA* genes, even though both hosts possess homologs of all four of these surface-receptor associated genes (Tables S2 and S3). On the phage side, S-RIM8 isolates coevolved with WH7803 or WH8101 had mutations in some of the same structural and tail proteins genes (but in different locations within the genes) (Fig. 3A; Table S7), indicating that S-RIM8 uses some of the same proteins to interact with different host-specific surface-receptors. Surprisingly, S-RIM8 isolates coevolved with WH7803 can infect all coevolved WH8101 strains, but not vice versa (Fig. 4).

In cases where pairwise coevolution with one partner did not generally alter interactions with another partner (light gray or white blocks in Fig. 4), completely different sets of surface-receptor genes were mutated in coevolved *Synechococcus* WH7803 isolates (Fig. 5C). For example, mutations in the *gal*E gene in WH7803 did not provide resistance to the ancestral or any of the coevolved S-RIM4 or S-RIM8 isolates. Likewise, mutations in receptor-related genes in ISL1 of WH7803 did not provide resistance to the ancestral S-CAM7 strains or any of the coevolved S-CAM7 isolates. These results suggest that the two S-CAM7 strains and S-RIM4/S-RIM8 are interacting with different sets of cell-surface receptors on WH7803 cells.

Beyond mutations providing extracellular resistance, mutations in genes likely involved in intracellular interactions often led to broad resistance across the PBIN (Fig. 5D). Two WH7803 isolates coevolved with S-RIM4 (7803RIM4B187-01 and 7803RIM4B187-06) were resistant to all ancestral and coevolved isolates of 1-S-CAM7, 2-S-CAM7, S-RIM4, and S-RIM8 as well as S-RIM44 and the two S-RIP4 strains (Figs. 2A and 4). These two WH7803 isolates possess a unique 7-nucleotide insertion leading to a frameshift in a serralysin-like metalloprotease (SynWH7803_0122). Recently, novel antiphage defense systems based on proteases have been discovered [44,51]. In another example, a frameshift mutation in a ribosome biogenesis GTPase (thiamine phosphate phosphatase) in WH7803 isolate (1-7803CAM7A19-03) coevolved with 1-S-CAM7 became resistant to all S-RIM4 isolates as well as most S-RIM8 isolates. Finally, some WH7803 isolates coevolved with 1-S-CAM7 possess a mutation in the upstream of a conserved hypothetical protein. This mutation provides resistance to some 1-S-CAM7 and 2-S-CAM7 phages as well as several S-RIM4 coevolved isolates (outlined in red, Fig. 4). While phage-resistance arising during pairwise coevolution was generally limited in its effect to one other phage partner, these particular mutations in genes that are likely involved in intracellular interactions allowed for extensive resistance across the infection network (Fig. 5D), suggesting that some intracellular infection processes are shared among the diverse phages used in this study. Notably, our results differ from other experimental evolution laboratory studies where extracellular receptor-mediated resistance in marine Flavobacteria [52] and *Salmonella enterica* [39] provided broad resistance against multiple phages while intracellular resistance provided much narrower resistance against a single phage.

## Conclusion

Here we show that pairwise coevolution influences PBINs both through changes in cell-surface as well as intracellular interactions, altering interactions among other members of the community. As hypothesized, the impact of coevolution on the interaction network depended on the degree to which interaction genes were shared among other phages and/or hosts. Overall, we found that the consequences of pairwise coevolution for interactions with other partners fell into four major categories (Fig. 5). Interactions were altered across pairs if phages interacted with the same set of cell surface receptor genes and largely modular if phages utilized different sets of host surface receptors. Non-reciprocal interactions were observed when phages utilized some of the same tail fiber genes to absorb to different sets of cell-surface receptors on host A than on host B. Here, coevolved phage isolates from one pairwise interaction could infect most or all coevolved host isolates from another pair, but not vice versa. In a few cases, we also observed host mutations that conferred broad resistance to many phage strains, regardless of the surface receptors used for adsorption, suggesting novel intracellular defense systems. Our results suggest that the structure of interaction networks from complex natural communities could be used to infer ongoing antagonistic coevolution and the types of genetic interactions underlying phage-host interactions. Further studies, however, are needed to better understand the mechanisms of intercellular phage defense systems, particularly those that provide broad resistance against diverse phages and thus can have a large impact on PBINs.

## Acknowledgments

We thank Brooke Fenderson, Caleb Milford, Alex Veglia, and Jonathan Abebe for assistance in the lab and Rich Puxty for input on the project.

## Data availability

All sequence data are available under BioProject ID PRJNA657566 and PRJNA1538709.

