## Supplemental Methods, Figure, Tables for "Genomic signatures of coevolving phage-bacteria interaction networks"

### Supplementary Materials and Methods

#### *Synechococcus* and cyanophage strains and growth conditions

Marine *Synechococcus* sp. WH7803 and *Synechococcus* sp. WH8101 were originally obtained from the Woods Hole Collection of Cyanobacteria (Woods Hole Oceanographic Institute, Woods Hole, MA) (Table S1) [1]. *Synechococcus* sp. WH7803 and WH8101 fall into different clades (V and VIII respectively) and differ in many physiological characteristics [2,3]. These *Synechococcus* cultures were not axenic and contained heterotrophic bacteria. Prior to the start of each experiment, a single colony of *Synechococcus* was isolated as previously described [4]. Briefly, cells were serially diluted, combined with 30 ml of SN soft agar, and grown on petri dishes in a growth chamber with 25 μmol photons $m^{-2}$ $s^{-1}$at 23°C with diurnal cycles (14 h of light). After single colonies were visible, individual colonies were removed from the soft agar plates and regrown in SN medium in an Erlenmeyer flask in the growth chamber.

Bacteriophage isolates used in this study were isolated from the coastal waters of Rhode Island or California as previously described [5-7] (Table S1). After initial isolation, filter-sterilized (0.22 μm) phage lysates were stored at 4°C. All phage isolates were plaque-purified again before initiation of evolution/coevolution experiments. This was accomplished by plating serial dilutions of viral lysates and concentrated host cells in soft agar on SN agar plates as previously described [8]. Viral plaques were picked with a toothpick and regrown on the ancestral host culture.

The four T4-like (myovirus) cyanophage strains (S-CAM7, S-RIM8, S-RIM4 and S-RIM44) are genetically diverse (Fig. S1). To examine differences in evolution/coevolution among very closely related isolates, two isolates belonging to S-CAM7 and two isolates belonging to S-RIP4 were used in this study. S-CAM7 0809CC03 and 0910SB42 have an overall genome ANI of 95%. S-RIP4a and S-RIP4b have a genome ANI of 88% and differ in a region of their genome that has been associated with tail fiber proteins [9].

**Coevolution and evolution experimental design**

Coevolution experiments were conducted in replicated semi-continuous batch culture for the S-CAM7 phage isolates and continuous culture in chemostats for the S-RIM and S-RIP phage isolates. For semi-continuous cultures, the ancestral host was inoculated in AN medium at $OD_{750}$ of 0.2 then grown over night. The next day the host was inoculated with the ancestral phage isolate in 20-mL of AN media at a concentration of $3x10^5$ phage/ml. Each subsequent week for a total of 19 weeks, the previous week's culture was used as an inoculum in a 50:50 substitution with fresh media into a new flask. Flasks were incubated at 25 μmol photons $m^{-2}$ $s^{-1}$ at 25°C on a 14:10 light-dark cycle. Coevolution experiments in continuous culture were conducted as previously described [10]. In brief, either an artificial seawater medium or SN medium was supplied to replicated 35-mL chemostats at a dilution of ~1.02/d. The contents were stirred with magnetic stir bars and maintained at 23.0 C on a 14:10 light:dark cycle at 10 μmol photons $m^{-2}$ $s^{-1}$.

In addition to the coevolution experiments, each viral-host pair was also used in an 'evolution' experiment conducted in semi-continuous culture. In this treatment, the virus was reintroduced to susceptible hosts at each transfer, allowing the virus to evolve and adapt to the ancestral host, but not to cells that had evolved resistance. For the evolution of S-CAM7, replicate 20 mL cultures of ancestral host and phage lysate were established as in the coevolution experiments. Each week 10-mL of the resulting phage lysate was filtered (Steriflip 0.2 μm) to remove any remaining host cells and then concentrated 10-fold by centrifugal filtration (Amicon Ultra-4 30kDa, 4500 x g, 4 min). This concentrated lysate was used to infect 10-mL of naïve host cells in AN medium. Viral lysates were transferred 19 times. For the 'evolution' experiments of

S-RIM4, S-RIM8 and S-RIP4, replicate 35-mL cultures of ancestral cells in SN media were inoculated with ancestral phage lysates. Every five to eight days (based on the rate of cell lysis), phage lysates were filtered (Corning Tube Top Vacuum Filter System, 0.22 μm) and 5-mL of filtered lysate was added to new flasks with 10-mL of naïve host cells and 20-mL SN media. For the S-RIP4 evolution experiment, starting with transfer 15, and every subsequent transfer, only 2.5-mL of filtered lysate was used to inoculate naïve host cells. S-RIM4 and S-RIM8 were transferred 25 times in evolution experiments using *Synechococcus* WH7803 and WH8101 as hosts. S-RIP4 isolates on *Synechococcus* WH7803 were transferred 24 times (Table 1). For the 'host evolution control' in semi-continuous culture, the ancestral host was serially transferred 50:50 into fresh media each week. For the continuous culture experiments, the 'host evolution control' consisted of replicated chemostats containing cells, but no virus.

**Isolation and genomic sequencing of *Synechococcus* and cyanophages**

Single colonies of *Synechococcus* were isolated from each of the coevolution and evolution experiments on soft agar plates and phages from the experiments were plaque-purified on the ancestral host as previously described [4,7,8]. Single *Synechococcus* colonies were picked and grown in SN as described above. Viral plaques were grown in the ancestral host strains. Genomic DNA from coevolution and evolution experiments with S-CAM7 isolates and *Synechococcus* strain WH7803 was extracted by following the protocol of the Quick gDNA Miniprep kit (D3006, ZymoResearch, Santa Ana, CA, USA) after concentrating samples from 2-4 ml to approximately 500 ul with an Amicon Ultra 30kDa cut off column. Libraries for whole genome sequencing were generated using the Nextera XT kit (Illumina) following the protocol provided with the kit. The libraries for each sample were then pooled and paired end sequenced (2x300bp) on an Illumina MiSeq at the UC Davis DNA core facility. Genomic DNA from the S-RIM and S-RIP phages was isolated using Promega Wizard DNA purification resin and spin columns as previously described (Henn et al., 2010). The PowerWater DNA isolation kit (MoBio Laboratories) or DNeasy PowerWater Kit (Qiagen) was used to isolate DNA from S-RIM and S-RIP phage communities as well as *Synechococcus* isolates and communities. Genomic DNA library preparation and Illumina MiSeq sequencing using a 500-cycle reagent kit V.2 (2x250bp) was performed at the University of Rhode Island Genomics and Sequencing Center or at the University of Delaware DNA Sequencing and Genotyping Center [7,11].

**Genomic Analyses**

The sequences from the ancestral cell and phage isolates used to initiate each co/evolution experiment were used as references to identify genetic changes that occurred during the experiments (Table S1). For *Synechococcus* strain WH7803, GenBank accession CT971583 was used as the reference sequence. When ancestral WH7803 isolates from each coevolution or evolution experiment were mapped to this sequence, they differed by 51 to 57 nucleotides from the reference sequence with coverage ranging from 45-130X. These changes were also found to be present in the corresponding coevolved and evolved WH7803 isolates and thus were not considered in further analyses [10]. The ancestral isolate of *Synechococcus* strain WH8101 (GenBank accession NZ_CP035914) was fully sequenced [11] as were each of the ancestral phages (Table S1). Prior to mapping, Illumina MiSeq reads were trimmed for quality control in Geneious Prime® 2024 (Error Probability Limit 0.01, Maximum low quality bases 2, Maximum ambiguities 0, with the option to Trim both 3' and 5' ends). The trimmed reads were mapped back to the corresponding ancestral sequences using the Geneious Prime® mapper in the Map-to-Reference Tool (custom sensitivity, minimum overlap identity of 95%, with gaps allowed). SNPs and small insertions and deletions (indels) were also identified using Geneious Prime® (minimum coverage of 80 reads for phages and 10 reads for cells, minimum variant frequency of 0.1, maximum variant P-value $10^{-4}$, minimum strand-bias P-value $10^{-1}$ when bias is over 70%.) For host cell and phage isolates, only SNPs or indels with >80% of reads supporting the mutation were included in the final analyses. For cell and host communities, all SNPs/indels with >10% supporting reads were considered

**References Supplemental Methods**

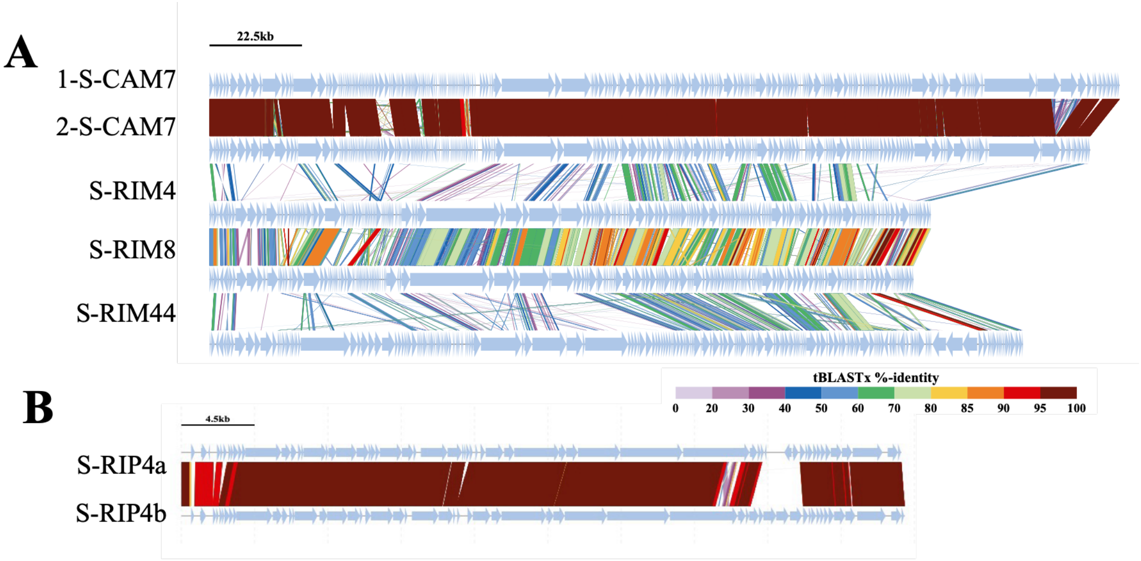


**Figure S1.** Genome alignments based on amino acids similarity for T4-like phage (A) and T7-like phage (B). Alignments were constructed using DiGAlign [12].

Table S1 *Synechococcus* and cyanophage strains used in this study

| Host/Phage Isolates | Classification | Isolation date | Isolation host | Location | Genome Size | GenBank Acession# | Reference |
|---|---|---|---|---|---|---|---|
| *Synechococcus* WH7803 | Cyanobacteria | 1978 | na | Atlantic Ocean | 2366980 | CT971583 | Waterbury et al 1986 |
| *Synechococcus* WH8101 | Cyanobacteria | 1981 | na | Woods Hole, MA | 2630292 | NZ_CP035914 | Waterbury et al 1986 |
| 1-S-CAM7_0809CC03 | T4-like cyanophage | August 2009 | Syn-WH7803 | Southern CA | 221362 | MT586120 | Crummet et al 2016 |
| 2-S-CAM7_0910SB42 | T4-like cyanophage | Sept 2009 | Syn-WH7803 | Southern CA | 214214 | KU686213 | Crummet et al 2016 |
| S-RIM8_RW_22_0300 | T4-like cyanophage | March 2000 | Syn-WH7803 | Narragansett Bay RI | 171216 | KX349289 | Marston & Martiny 2016 |
| S-RIM4_RW_11_0999 | T4-like cyanophage | Sept 1999 | Syn-WH8012 | Narragansett Bay RI | 175462 | MK493321 | This study |
| S-RIM44_Np_20_0711 | T4-like cyanophage | July 2011 | Syn-WH7803 | Narragansett Bay RI | 197868 | KX349293 | Marston & Martiny 2016 |
| S-RIP4a_RW_07_0909 | T7-like cyanophage | Sept 2009 | Syn-WH7803 | Narragansett Bay RI | 44194 | PV872382 | This study |
| S-RIP4b_RW_20_0813 | T7-like cyanophage | August 2013 | Syn-WH7803 | Narragansett Bay RI | 44406 | PV872383 | This study |

Table S2 Loci with mutations in coevolved *Synechococcus* WH7803 isolates

| Cyanorak locus tag | Genomic Island | Core Gene | Homolog in WH18101 | operon number | CLOG | Manually Edited | Product Description | *Synechococcus* WH7803 coevolved with [a] | | | | | | | | | | | | | | |
|---|---|---|---|---|---|---|---|---|---|---|---|---|---|---|---|---|---|---|---|---|---|---|
| | | | | | | | | 1-S-CAM7 | | | | 2-S-CAM7 | | S-RIM4 | | S-RIM8 | | | S-RIP4a | | S-RIP4b | |
| | | | | | | | | A | B | C | D | B | D | A | B | A | C | D | F | G | F | G |
| | | | | | | | | 4 | 4 | 4 | 4 | 4 | 4 | 5 | 4 | 3 | 5 | 5 | 1 | 3 | 3 | 3 |
| SynWH7803_0025 | no | yes | yes | OP48 | CK_00000349 | yes | ribosome biogenesis GTPase / thiamine phosphate phosphatase | 1 | | | | | | | | | | | | | | |
| SynWH7803_0064 | no | yes | yes | none | CK_00000325 | yes | ATP-dependent Clp protease ATP-binding subunit, ClpX | | | | | | | | | | 1 | | | | | |
| SynWH7803_0076 | no | yes | yes | OP82 | CK_00000319 | yes | tRNA(Ile)-lysidine synthetase TilS | 1 | | | | | | | | | | | | | | |
| SynWH7803_0092 | GI01 | no | yes | OP91 | CK_00000090 | yes | galE UDP-glucose 4-epimerase | | 4 | | | | | | | | | | | | | |
| IGR 5'-SynWH7803_0102 | GI01 | | | OP96 | na | na | | | | | | | | | 4 | 2 | 3 | 2 | 1 | 3 | 1 | 3 |
| SynWH7803_0102 | GI01 | no | yes | OP96 | CK_00001513 | yes | rfbA, glucose-1-phosphate thymidylyltransferase | | | | | | | | | 1 | 1 | 3 | | | | |
| SynWH7803_0104 | GI01 | no | yes | OP96 | CK_00042324 | yes | rfbD, dTDP-4-dehydrorhamnose reductase | | | | | | | | | | 1 | | | | | |
| SynWH7803_0105 | GI01 | no | no | OP96 | CK_00001954 | yes | wcaJ, undecaprenyl-phosphate glucose phosphotransferase | | | | | | | 4 | | | | | | | 2 | |
| SynWH7803_0108 | GI01 | no | no | none | CK_00048005 | no | hypothetical protein | | | | | | | 1 | | | | | | | | |
| SynWH7803_0122 | GI01 | no | no | none | CK_00039995 | yes | serralysin-like metalloprotease | | | | | | | | 2 | | | | | | | |
| SynWH7803_0133 | GI01 | no | no | OP112 | CK_00007532 | no | rhamnan synthesis F family protein | | | | | | | 1 | 1 | | | | | | | |
| IGR 5'-SynWH7803_0133 | GI01 | | | none | na | | | | | | | | | 1 | | | | | | | | |
| SynWH7803_0134 | GI01 | no | no | OP113 | CK_00057572 | yes | hypothetical protein | | | | | | | 1 | | | | | | | | |
| SynWH7803_0137 | GI01 | no | no | OP113 | CK_00002497 | yes | ABC-type lipopolysaccharide transport system, permease component | | | | | | | | 1 | | | | | | | |
| SynWH7803_0139 | GI01 | no | no | OP114 | CK_00002970 | no | possible alpha-glycosyltransferase, family 4 | | | | | | | | | 1 | | | | | | |
| SynWH7803_0218 | no | no | yes | none | CK_00001335 | no | beta-glycosyltransferase, family 2 | | | | | | | | | | | | | | 1 | 2 |
| SynWH7803_0219 | no | no | yes | none | CK_00001544 | no | hypothetical protein | | | | | | | | | | | | 1 | | | |
| SynWH7803_0242 | no | no | yes | OP169 | CK_00000090 | yes | galE, UDP-glucose 4-epimerase | 1 | 4 | 1 | | 4 | 1 | | | | | | | | | |
| IGR 5' SynWH7803_0242 | no | | | OP169 | na | | | | | 1 | 4 | | | | | | | | | | | |
| SynWH7803_0256 | no | yes | yes | OP176 | CK_00008067 | no | ABC transporter family protein | | | 2 | | | | | | | | | | | | |
| SynWH7803_0379 | no | no | yes | OP249 | CK_00000494 | yes | elongation factor EF-Tu | | | | | | | | | 1 | | | | | | |
| SynWH7803_0771 | no | yes | yes | OP485 | CK_00000932 | yes | prlC, oligopeptidase A | | | | | | | | 1 | | | | | | | |
| SynWH7803_0948 | GI05 | no | no | none | CK_00003002 | yes | cupredoxin-like protein | | | | | | | | | 1 | | | | | | |
| SynWH7803_1075 | no | no | yes | OP694 | CK_00001815 | yes | Arsenite efflux pump ACR3 related permease, arsB | | | | | | | | 1 | | | | | | | |
| SynWH7803_1116 | no | yes | yes | OP720 | CK_00008087 | yes | possible photosystem II D1 repair protein FtsH3 | | | | | | | | | | 1 | | | | | |
| SynWH7803_1265 | no | yes | yes | none | CK_00001458 | no | hypothetical protein | | | | | | | | | | | 1 | | | | |
| SynWH7803_1342 | no | yes | yes | OP864 | CK_00000899 | no | 50S ribosome-binding GTPase family protein | | | | | | | 1 | | | | | | | | |
| SynWH7803_1391 | no | no | yes | OP898 | CK_00002870 | yes | RecF-like Rad50/SbcC-type AAA domain-containing protein | 1 | | | | | | | | | | | | | | |
| SynWH7803_1435 | no | yes | yes | OP928 | CK_00000965 | yes | triosephosphate isomerase | | | | | | | | | | | | 1 | | | |
| SynWH7803_1446 | no | yes | yes | none | CK_00001292 | no | hypothetical protein | | | | | | | | | | | 1 | | | | |
| SynWH7803_1499 | no | no | no | none | CK_00057406 | yes | hypothetical protein | | | | | | | | 1 | | | | | | | |
| 5' SynWH7803_1505 and SynWH7803_1506 | no | | | none | na | na | | 3 | | 2 | | | | | | | | | | | | |
| SynWH7803_1507 | no | yes | yes | OP973 | CK_00000748 | yes | exodeoxyribonuclease V (RecBCD complex), gamma subunit | | | | | | | | | | | 2 | | | | |
| SynWH7803_1797 | no | no | yes | OP1145 | CK_00002031 | yes | uncharacterized conserved membrane protein | | | | | | 1 | | | | | | | | | |
| 5' SynWH7803_1798 | no | | | none | na | na | | | | | | | 1 | | | | | | | | | |
| 5' SynWH7803_2034 | no | | | none | na | na | | | | | | | | 1 | | | | | | | | |
| SynWH7803_2069 | GI13 | no | no | none | CK_00057421 | yes | carbohydrate-binding protein (CBM2) | | | | | | | | | | | 1 | | | | |
| SynWH7803_2399 | no | no | yes | OP1501 | CK_00001684 | no | conserved hypothetical protein | | | | 3 | | | | | | | | | | | |
| SynWH7803_2428 | no | yes | yes | OP1517 | CK_00046122 | yes | DNA ligase, NAD-dependent, ligA | | | | 4 | 1 | 3 | | | | | | | | | |
| SynWH7803_2466 | no | no | yes | OP1534 | CK_00056759 | yes | ABC-type multidrug transport system, ATPase and permease components | | | | | | | | | 1 | | | | | | |

[a] Letters in the top row are the replicate designation. The second row is the number of isolates sequenced in each replicate. The number in the columns below represents the number of isolates within the replicate that possessed at least one mutation in the gene/region.

Table S3  Loci with mutations in coevolved *Synechococcus* WH8101 isolates

| Cyanorak locus tag | Genomic Island | Core Gene | Homolog in WH7803 | CLOG | Manually Edited | Product Description | *Synechococcus* WH8101 coevolved with [a] | | |
|---|---|---|---|---|---|---|---|---|---|
| | | | | | | | S-RIM8 | | S-RIM44 |
| | | | | | | | A | B | A |
| | | | | | | | 7 | 6 | 2 |
| SynWH8101_00118 | no | no | yes | CK_00000025 | yes | alpha/beta hydrolase fold family protein | | | 1 |
| SynWH8101_00120 | no | no | yes | CK_00009125 | yes | RND family multidrug efflux transporter, MMPL family | 1 | | |
| SynWH8101_00148 | no | no | yes | CK_00001338 | yes | phosphotransferase enzyme family protein | | 5 | 1 |
| SynWH8101_00204 | GI03 | no | yes | CK_00001332 | yes | kdsD, arabinose-5-phosphate isomerase | 4 | | |
| SynWH8101_00216 | GI03 | no | yes | CK_00001666 | yes | P-loop containing nucleoside triphosphate hydrolase | 4 | 6 | |
| SynWH8101_00269 | GI05 | no | no | CK_00050082 | yes | outer membrane autotransporter barrel domain-containing protein | | | 1 |
| SynWH8101_00299 | no | yes | yes | CK_00009007 | yes | degT, putative pleiotropic regulatory protein | | 4 | |
| SynWH8101_00493 | no | yes | yes | CK_00000537 | no | haloacid dehalogenase-like hydrolase family protein | | 1 | |
| SynWH8101_00631 | no | yes | yes | CK_00001043 | yes | conserved hypothetical protein YlmE | 4 | | |
| SynWH8101_00843 | no | yes | yes | CK_00000042 | yes | photosystem II D2 protein | 1 | | |
| SynWH8101_01139 | no | yes | yes | CK_00008111 | no | uncharacterized membrane protein | 2 | | |
| SynWH8101_01325 | GI14 | no | yes | CK_00057091 | yes | DNA primase catalytic core, N-terminal domain protein | | | 1 |
| SynWH8101_01387 | no | no | yes | CK_00001285 | yes | homoserine O-succinyltransferase | | 5 | 1 |
| SynWH8101_01646 | no | no | yes | CK_00001469 | yes | lipopolysaccharide export system, ATPase component, lptA | | 4 | |
| SynWH8101_01666 | no | yes | yes | CK_00000896 | yes | metallopeptidase, M16 family | | 1 | |
| SynWH8101_02225 | no | yes | yes | CK_00001581 | yes | peptidase M41 family | | | 1 |
| SynWH8101_02290 | no | yes | yes | CK_00001410 | yes | conserved hypothetical protein | 4 | | |
| SynWH8101_02368 | no | yes | yes | CK_00001641 | yes | circadian clock protein kaiA | 1 | | |
| SynWH8101_02462 | no | yes | yes | CK_00001058 | yes | high light inducible protein | 1 | | |
| 5' SynWH8101_02478 | no | yes | yes | na | yes | ribosome maturation factor RimP | 1 | | |
| SynWH8101_02502 | no | no | yes | CK_00004865 | no | NHPM bacteriocin system secretion protein, HlyD family | 1 | | |
| SynWH8101_02536 | no | yes | yes | CK_00000187 | yes | peptide deformylase | 2 | | |
| SynWH8101_02794 | no | yes | yes | CK_00000414 | yes | alanyl-tRNA synthetase | | | 1 |
| SynWH8101_02830 | no | yes | yes | CK_00000401 | yes | valyl-tRNA synthetase | 2 | | |
| SynWH8101_02892 | no | no | yes | CK_00001554 | yes | possible chelatase | | | 1 |

[a] Letters in the top row are the replicate designation. The second row is the number of isolates sequenced in each replicate. The number in the columns below represents the number of isolates within the replicate that possessed at least one mutation in the gene/region.

Table S4 Loci with mutations in coevolved *Synechococcus* WH7803 and WH8101 by function

| Cyanorak locus tag | Genomic Island | Gene Product Description | WH7803 coevolved with [a] | | | | | | | | | | | | | | | WH8101 with | |
|---|---|---|---|---|---|---|---|---|---|---|---|---|---|---|---|---|---|---|---|
| | | | 1-7803CAM7 | | | | 2-7803CAM7 | | 7803RIM4 | | 7803RIM8 | | | 7803RIP4a | | 7803RIP4b | | 8101RIM8 | | 8101RIM44 |
| | | | A | B | C | D | B | D | A | B | A | C | D | F | G | F | G | A | B | A |
| | | | 4 | 4 | 4 | 4 | 4 | 4 | 5 | 4 | 3 | 5 | 5 | 1 | 3 | 3 | 3 | 7 | 6 | 2 |
| **Surface polysaccharides, membrane biosynthesis and transport** | | | | | | | | | | | | | | | | | | | | |
| SynWH7803_0092 | GI01 | galE, UDP-glucose 4-epimerase | | 4 [b] | | | | | | | | | | | | | | | | |
| IGR 5'-SynWH7803_0102 | GI01 | * 5' rfbA | | | | | | | | 4 | 2 | 3 | 2 | 1 | 3 | 1 | 3 | | | |
| SynWH7803_0102 | GI01 | * rfbA, glucose-1-phosphate thymidylyltransferase | | | | | | | | | 1 | 1 | 3 | | | | | | | |
| SynWH7803_0104 | GI01 | * rfbD, dTDP-4-dehydrorhamnose reductase | | | | | | | | | | 1 | | | | | | | | |
| SynWH7803_0105 | GI01 | * wcaJ, undecaprenyl-phosphate glucose phosphotransferase | | | | | | | 4 | | | | | | | 2 | | | | |
| SynWH7803_0133 | GI01 | rhamnan synthesis F family protein | | | | | | | 1 | 1 | | | | | | | | | | |
| SynWH7803_0137 | GI01 | ABC-type lipopolysaccharide transport system | | | | | | | | 1 | | | | | | | | | | |
| SynWH7803_0139 | GI01 | possible alpha-glycosyltransferase, family 4 | | | | | | | | | 1 | | | | | | | | | |
| SynWH7803_0218 | no | beta-glycosyltransferase, family 2 | | | | | | | | | | | | | | 1 | 2 | | | |
| SynWH7803_0242 | no | galE, UDP-glucose 4-epimerase | 1 | 4 | 1 | | 4 | 1 | | | | | | | | | | | | |
| IGR 5' SynWH7803_0242 | no | 5' galE | | | 1 | 4 | | | | | | | | | | | | | | |
| SynWH7803_1435 | no | triosephosphate isomerase | | | | | | | | | | | | 1 | | | | | | |
| SynWH7803_1797 | no | uncharacterized conserved membrane protein | | | | | | 1 | | | | | | | | | | | | |
| SynWH7803_2069 | GI13 | carbohydrate-binding protein (CBM2) | | | | | | | | | | | 1 | | | | | | | |
| SynWH8101_00204 | GI03 | kdsD, arabinose-5-phosphate isomerase | | | | | | | | | | | | | | | | 4 | | |
| SynWH8101_00269 | GI05 | outer membrane autotransporter | | | | | | | | | | | | | | | | | | 1 |
| SynWH8101_00299 | no | degT, putative pleiotropic regulatory protein | | | | | | | | | | | | | | | | | 4 | |
| SynWH8101_01139 | no | uncharacterized membrane protein | | | | | | | | | | | | | | | | 2 | | |
| SynWH8101_01646 | no | lptA, lipopolysaccharide export system, ATPase component, | | | | | | | | | | | | | | | | | 4 | |
| **Translation** | | | | | | | | | | | | | | | | | | | | |
| SynWH7803_0025 | no | ribosome biogenesis GTPase / thiamine phosphate phosphatase | 1 | | | | | | | | | | | | | | | | | |
| SynWH7803_1342 | no | 50S ribosome-binding GTPase family protein | | | | | | | 1 | | | | | | | | | | | |
| SynWH7803_0076 | no | tRNA(Ile)-lysidine synthetase TilS | 1 | | | | | | | | | | | | | | | | | |
| SynWH7803_0379 | no | elongation factor EF-Tu | | | | | | | | | 1 | | | | | | | | | |
| 5' SynWH8101_02478 | no | ribosome maturation factor RimP | | | | | | | | | | | | | | | | 1 | | |
| SynWH8101_02794 | no | alanyl-tRNA synthetase | | | | | | | | | | | | | | | | | | 1 |
| SynWH8101_02830 | no | valyl-tRNA synthetase | | | | | | | | | | | | | | | | 2 | | |
| **Protein folding, stability** | | | | | | | | | | | | | | | | | | | | |
| SynWH7803_0064 | no | ATP-dependent Clp protease ATP-binding subunit, ClpX | | | | | | | | | | 1 | | | | | | | | |
| SynWH7803_0122 | GI01 | serralysin-like metalloprotease | | | | | | | | 2 | | | | | | | | | | |
| SynWH7803_0771 | no | prlC, oligopeptidase A | | | | | | | | 1 | | | | | | | | | | |
| SynWH7803_2466 | no | ABC-type multidrug transport system, ATPase and permease | | | | | | | | | 1 | | | | | | | | | |
| SynWH8101_01387 | no | homoserine O-succinyltransferase | | | | | | | | | | | | | | | | | 5 | 1 |
| SynWH8101_01666 | no | metallopeptidase, M16 family | | | | | | | | | | | | | | | | | 1 | |
| SynWH8101_02225 | no | peptidase M41 family | | | | | | | | | | | | | | | | | | 1 |
| SynWH8101_02536 | no | peptide deformylase | | | | | | | | | | | | | | | | 2 | | |
| **DNA replication, recombination, repair** | | | | | | | | | | | | | | | | | | | | |
| SynWH7803_1391 | no | RecF-like Rad50/SbcC-type AAA domain-containing protein | 1 | | | | | | | | | | | | | | | | | |
| SynWH7803_1507 | no | exodeoxyribonuclease V (RecBCD complex), gamma subunit | | | | | | | | | | | 2 | | | | | | | |
| SynWH7803_2428 | no | DNA ligase, NAD-dependent, ligA | | | | 4 | 1 | 3 | | | | | | | | | | | | |
| SynWH8101_01325 | GI14 | DNA primase catalytic core, N-terminal domain protein | | | | | | | | | | | | | | | | | | 1 |
| **Efflux pumps, secretion** | | | | | | | | | | | | | | | | | | | | |
| SynWH7803_0256 | no | ABC transporter family protein | | | 2 | | | | | | | | | | | | | | | |
| SynWH7803_1075 | no | Arsenite efflux pump ACR3 related permease, arsB | | | | | | | | 1 | | | | | | | | | | |
| SynWH8101_02502 | no | NHPM bacteriocin system secretion protein, HlyD family | | | | | | | | | | | | | | | | 1 | | |

[a] Letters in the top row are the replicate designation. The second row is the number of isolates sequenced in each replicate. The number in the columns below represents the number of isolates within the replicate that possessed at least one mutation in the gene/region.

Table S5  Loci with mutations in S-CAM7 isolates and communities coevolved or evolved with *Synechococcus* WH7803

| 1-S-CAM7 locus tag | 2-S-CAM7 locus tag | product | ISL [a] | PHROG | PHROG Annotation | Functional Category | 1-S-CAM7 Coevolved WH7803 | | 1-S-CAM7 Evolved WH7803 | | | | | | 2-S-CAM7 Coevolved WH7803 | | 2-S-CAM7 Evolved WH7803 | | | |
|---|---|---|---|---|---|---|---|---|---|---|---|---|---|---|---|---|---|---|---|---|
| | | | | | | | **A** [b] | **C** | **E** | **F** | **1** | **2** | **3** | **4** | **B** | **D** | **1** | **2** | **3** | **4** |
| | | | | | | | **4** | **3** | **3** | **3** | **C** | **C** | **C** | **C** | **4** | **4** | **C** | **C** | **C** | **C** |
| CC030809_00013 # | S420910_014- 015 | hypothetical protein | | 28263 | tail fiber protein | tail | 1 | 1 | | | | | | | 4 | 3 | X | X | X | X |
| CC030809_00018 | S420910_020 | hypothetical protein | | 529 | tail fiber protein | tail | | 3 | 3 | 3 | X | X | X | X | | | | | | |
| CC030809_00023 | S420910_024 | hypothetical protein | TF-BP | 16 | DNA helicase | nucleotide metabolism | | | | 1 | | | | | | | | | | |
| 5'CC030809_00028# | S420910_029 - 033 | all hypothetical proteins | TF-BP | 5 ORF insertion | | | 1 | 3 | 3 | 3 | X | X | X | X | | | | | | |
| not present | S420910_030 | hypothetical protein | TF-BP | 3434 | hpothetical | unknown | | | | | | | | | **1** | | | | | |
| CC030809_00050 | not present | hypothetical protein | TF-BP | none | hypothetical | unknown | | **1** | | | | | | | | | | | | |
| CC030809_00056 | S420910_057 | hypothetical protein | TF-BP | 21201* | DNA methylase | other | | | | | | | | | | **1** | | | | |
| CC030809_00058 | S420910_059 | hypothetical protein | TF-BP | 3,041* | cytidyltransferase | other | | | | 1 | | | | | | | | | | |
| CC030809_00072 - 00074 | S420910_072-074 | hypothetical proteins | TF-BP | 3435; 28768; 522 | all hypothetical | unknown | **1** | | | | | | | | | | | | | |
| CC030809_00076 | S420910_076 | hypothetical protein | TF-BP | 21201* | DNA methylase | unknown | | | | 1 | | | | | 2 | | | | | |
| 5' CC030809_00078 | 5' S420910_077 | IGR | TF-BP | | | | | | | | | X | X | | | | | | | |
| CC030809_00080 | S420910_079 | hypothetical protein | TF-BP | 3508 | hypothetical | unknown | | | | | | | | | | 1 | | | | |
| CC030809_00086 | S420910_086 | hypothetical protein | TF-BP | none | | unknown | | 1 | | 3 | | | | | | | | | | |
| CC030809_00087 | S420910_088 | hypothetical protein | TF-BP | 33546 | hypothetical | unknown | | | 3 | 3 | X | X | X | X | | | X | X | X | X |
| 5' CC030809_00088 | S420910_089 | IGR | TF-BP | | | | **1** | **3** | | | | | | | | | | | | |
| CC030809_00098 - 00105 | S420910_101 - 108 | hypothetical proteins | TF-BP | 3146; 31379; 3367; 35291; 871; 28075; 32260; 3298 | all hypothetical | unknown | | 1 | | | | | | | | | | | | |
| CC030809_00105 | S420910_108 | hypothetical protein | TF-BP | 3298 | hypothetical | unknown | | | 2 | | | | | | | | | | | |
| 5'CC030809_00109 | 5'S420910_112 | IGR | TF-BP | none | | | | | | 1 | | | | | | | | | | |
| CC030809_00114 | S420910_119 - 120 | hypothetical protein | TF-BP | 4492 | tail fiber | tail | 1 | | 3 | 3 | X | X | X | X | 1 | | X | X | X | X |
| CC030809_00115 | S420910_120 | hypothetical protein | TF-BP | 4492 | tail fiber | tail | | 1 | | | | | | | 1 | | X | X | X | X |
| CC030809_00116 | S420910_121 | hypothetical protein | TF-BP | 4492 | tail fiber | tail | | 3 | | | | X | | | | | | | | |
| CC030809_00122 | S420910_127 | hypothetical protein | | 2885 | baseplate wedge subunit | tail | | | | | X | | | X | | | | | | |
| CC030809_00148 | S420910_152 | RNA polymerase sigma factor for late transcription | | 234 | late sigma transcription factor | transcription regulation | | | | | | | | | | | | X | | |
| CC030809_00151 | S420910_155 | recombination-related endonuclease | | 77 | SbcC-like subunit of palindrome specific endonuclease | nucleotide metabolism | | | | | | | | | | | | X | X | |
| CC030809_00153 | S420910_157 | peptidase | | 459 | peptidase | other | 1 | | 3 | 1 | X | X | | X | | | X | X | X | X |
| CC030809_00156 | S420910_160 | CobS | | 249 | porphyrin biosynthesis | other | | | | | | | | | | 1 | | | | |
| CC030809_00162 | S420910_164 | hypothetical protein | | 18186* | baseplate wedge subunit | tail | | | | | | | | | 2 | 2 | X | X | X | X |
| CC030809_00163 | S420910_165 | hypothetical proteins | | 18186* | baseplate wedge subunit | tail | | **3** | | | | | | | | **2** | | | | |
| CC030809_00164 | S420910_166 | hypothetical protein | | 697* | structural protein | structural | | | | | | | | | **2** | | | | | |
| tRNA region loss of 4tRNA | tRNA region loss of **3 tRNAs** | | | | tRNA-Pro, **tRNA-Ser, tRNA-Thr, tRNA-Arg** | translation | 4 | 3 | | | | | | | 3 | 3 | X | X | X | X |
| tRNA | tRNA loss of 1 tRNA | | | | tRNA-ASN | translation | | | | | | | | | | | | | | X |
| CC030809_00237 | S420910_235 | hypothetical protein | | 28263 | tail fiber protein | tail | 1 | | | 1 | | | | | | 2 | | | | |
| CC030809_00238 | S420910_236 | hypothetical protein | | 28263* | tail fiber protein | tail | 1 | 2 | | | | | | | | | | | | |
| CC030809_00239 | S420910_237 | hypothetical proteins | | 2142* | tail fiber* | tail | 2 | 2 | | 1 | | | | | | | | | | |
| CC030809_00242 | S420910_242 | hypothetical protein | | 5073* | tail fiber* | tail | 2 | | 2 | 1 | | | | | | | | | | |
| CC030809_00243 | S420910_243 | hypothetical protein | | 22609* | tail fiber* | tail | 3 | | 3 | 2 | | | | | 1 | 1 | | | | |
| CC030809_00256 | S420910_251 | hypothetical protein | | 17885* | tail fiber* | tail | 1 | | | | | | | | | | | | | |
| CC030809_00258 | S420910_253 | hypothetical proteins | | 18186* | baseplate wedge subunit | tail | 1 | 3 | | | | | | | | | | | | |
| CC030809_00261 | S420910_254 | hypothetical protein | | 18186* | baseplate wedge subunit | tail | | | | | | | | | 1 | | | | | |
| CC030809_00268 | S420910_259 | hypothetical protein | | 2848 | hypothetical | unknown | 1 | | | | | | | | | | | | | |
| CC030809_00273 | S420910_263 | hypothetical protein | | 254 | late promoter transcriptional regulator | gene regulation | 1 | | | 1 | | | | | | 1 | | | | |

[a] Genomic Island. [b] Letters or numbers in top row are the replicate designation. The second row is the number of isolates sequenced in each replicate, 'C' indicates that the community lysate was sequenced. The number in the columns represents the number of isolates within the replicate that possessed at least one mutation in the gene/region, an "X" in the column indicates that at least one mutation was found in this gene/region in the community lysate sequence data. *PHROG based on phold annotations

Table S6 Loci with mutations in S-RIM4 isolates or communities coevolved or evolved with *Synechococcus* WH7803

| S-RIM4 locus_tag | Product | ISL [a] | PHROG | PHROG Annotation | Functional Category | Coevolved WH7803 | | Evolved WH7803 | |
|---|---|---|---|---|---|---|---|---|---|
| | | | | | | **A** [b] | **B** | **1** | **2** |
| | | | | | | 8 | 7 | C | C |
| RW110999_011 | hypothetical protein | | 790 | endolysin | lysis | | 3 | | |
| RW110999_012 | hypothetical protein | | 1200* | endolysin | lysis | | 1 | | |
| RW110999_025 | hypothetical protein | | 81 | 2OG-Fe(II) oxygenase | moron, auxiliary metabolic gene and host takeover | 7 | 3 | | |
| RW110999_031 | hypothetical protein | | 81 | 2OG-Fe(II) oxygenase | moron, auxiliary metabolic gene and host takeover | 7 | 6 | | |
| RW110999_039 | hypothetical protein | | 13910 | hypothetical protein | unknown function | 1 | | | |
| RW110999_045 | tail fiber protein | | 529 | tail fiber protein | tail | | | X | X |
| RW110999_047 | hypothetical protein | TF | 863 | tail fiber assembly | tail | 8 | 7 | | |
| RW110999_082 | hypothetical protein | TF | 573 | hypothetical protein | unknown function | | 2 | | |
| RW110999_094 | baseplate wedge subunit | | 219 | baseplate wedge subunit | tail | | 1 | | |
| RW110999_095 | structural protein | | 701 | virion structural protein | head and packaging | 2 | | | |
| RW110999_097 | hypothetical protein | | 708 | baseplate wedge subunit | tail | | 2 | | |
| RW110999_098 | baseplate wedge protein | | 438 | baseplate wedge subunit | tail | 2 | | | |
| RW110999_099 | hypothetical protein | | 18802 | virion structural protein | head and packaging | 3 | | X | X |
| RW110999_102 | structural protein | | 697 | structural protein | head and packaging | 3 | | | |
| RW110999_134 | major capsid protein | | 138 | major head protein | head and packaging | | 2 | | |
| RW110999_136 | 2OG-Fe(II) oxygenase family enzyme | | 61 | 2OG-Fe(II) oxygenase | moron; auxiliary metabolic gene and host takeover | | 1 | | |
| RW110999_138 | hypothetical protein | | 6464 | hypothetical protein | unknown function | | | X | X |
| RW110999_144 | hypothetical protein | | 1644 | minor coat protein | head and packaging | 1 | 1 | | |
| RW110999_162-169 | | ME | | | | 1 | | | |
| RW110999_166 | hypothetical protein | ME | 570 | hypothetical protein | unknown function | | | X | X |
| RW110999_169 | hypothetical protein | ME | 848 | hypothetical protein | unknown function | 2 | 7 | X | |
| RW110999_227 | hypothetical protein | | 417 | hypothetical protein | unknown function | | 1 | | |

[a] Genomic Island. [b] Letters or numbers in top row are the replicate designation. The second row is the number of isolates sequenced in each replicate, 'C' indicates that the community lysate was sequenced. The number in the columns represents the number of isolates within the replicate that possessed at least one mutation in the gene/region, an "X" in the column indicates that at least one mutation was found in this gene/region in the community lysate sequence data. *PHROG based on phold annotations

Table S7  Loci with mutations in S-RIM8 isolates and communities coevolved or evolved with *Synechococcus* WH7803 or WH8101

| S-RIM8 locus_tag | Product | ISL [a] | PHROG | PHROG Annotation | Functional Category | Coevolved WH7803 | | Evolved WH7803 | | | | Coevolved WH8101 | | Evolved WH8101 | | | |
|---|---|---|---|---|---|---|---|---|---|---|---|---|---|---|---|---|---|
| | | | | | | **C** [b] | **D** | **2** | **2** | **3** | **3** | **A** | **B** | **1** | **1** | **2** | **2** |
| | | | | | | 6 | 7 | 3 | C | 2 | C | 9 | 11 | 2 | C | 3 | C |
| RW220300_012 | hypothetical protein | | 1200 | endolysin | lysis | | **1** | | | | | | | | | | |
| RW220300_034 | structural protein | | 693 | tail fiber protein | tail | | **1** | | | | | | | | | | |
| RW220300_035 | tail fiber protein | | 529 | tail fiber protein | tail | | | **1** | | | | | **9** | | | | |
| RW220300_037 | hypothetical protein | TF-BP | 9851* | tail fiber protein | tail | **4** | **6** | **1** | **X** | **1** | **X** | **9** | **1** | **2** | **X** | **3** | **X** |
| RW220300_038 | hypothetical protein | TF-BP | 2504 | hypothetical protein | unknown function | | | | | | | **1** | | | | | |
| RW220300_039 | hypothetical protein | TF-BP | 81 | 2OG-Fe(II) oxygenase | auxiliary metabolic gene and host takeover | | | | | | | **3** | | | | | |
| RW220300_041 | hypothetical protein | TF-BP | 1464 | hypothetical protein | unknown function | | | | | | | | | | **X** | **3** | **X** |
| RW220300_068 | hypothetical protein | TF-BP | 2584 | hypothetical protein | unknown function | | | | | | | | | | | | **X** |
| 5'RW220300_077 | hypothetical protein | TF-BP | 5' 718 | hypothetical protein | unknown function | | | | | | | | | **2** | **X** | **1** | **X** |
| RW220300_078 | hypothetical protein | TF-BP | 707 | hypothetical protein | unknown function | | | | | | | | | | | **1** | |
| RW220300_086 | hypothetical protein | | 701 | virion structural protein | head and packaging | | | | | | | **8** | **9** | **2** | **X** | **3** | **X** |
| RW220300_088 | hypothetical protein | | 708 | baseplate wedge subunit | tail | | | | **X** | **2** | **X** | **5** | | | | | **X** |
| RW220300_089 | baseplate wedge subunit | | 438 | baseplate wedge subunit | tail | **6** | **7** | | | | | | **7** | **1** | | | |
| RW220300_090 | hypothetical protein | | 18802* | virion structural protein | head and packaging | | **1** | | | | | **6** | | | | | |
| RW220300_093 | structural protein | | 697 | structural protein | head and packaging | **1** | | | | | | | | | | | |
| RW220300_099 - 114 | minus 16 ORFs | gp16_gp17 | | | | | | | | | | | **9** | **2** | **X** | **2** | **X** |
| RW220300_122 | major capsid protein | | 138 | major head protein | head and packaging | **5** | **7** | **2** | **X** | **1** | **X** | | | | | | |
| RW220300_124 | hypothetical protein | | 6464 | hypothetical protein | unknown function | | **8** | **3** | **X** | **2** | **X** | | | | | | |
| RW220300_151 | translational repressor of early genes | | 242 | translation repressor | other | | | | | | | **1** | | | | | |
| RW220300_152 | hypothetical protein | | 2714 | hypothetical protein | unknown function | | | | | | | | | | **X** | **1** | **X** |
| RW220300_157 | hypothetical protein | | 570 | hypothetical protein | unknown function | **1** | | | | | | | **1** | | | | |
| RW220300_164 - 171 | minus 8 ORFs | ME | | | | **2** | | | | | | | | | | | |
| RW220300_166 | hypothetical protein | ME | 848 | hypothetical protein | unknown function | **2** | **5** | **1** | **X** | **1** | **X** | **1** | **1** | | | | |
| RW220300_186 | hypothetical protein | | 1784 | hypothetical protein | unknown function | | | | | | | | | **1** | | | |
| RW220300_186-188 | hypothetical proteins | | delection | | | **1** | | | | | | | | | | | |
| RW220300_188 | hypothetical protein | | 613 | head protein | head and packaging | | | | | | | **1** | **9** | | | | |
| RW220300_191 | hypothetical protein | | 451 | hypothetical protein | unknown function | | **1** | | | | | | | | | | |
| RW220300_216 | photosystem II protein D | psbA-psbD | 182 | photosystem II D1 | auxiliary metabolic gene and host takeover | | | **1** | **X** | | | | | | | | |
| RW220300_218 | hypothetical protein | | 1033* | ParB-like partition protein | DNA, RNA and nucleotide metabolism | | **1** | | | | | | | | | | |

[a] Genomic Island. [b] Letters or numbers in top row are the replicate designation. The second row is the number of isolates sequenced in each replicate, 'C' indicates that the community lysate was sequenced. The number in the columns represents the number of isolates within the replicate that possessed at least one mutation in the gene/region, an "X" in the column indicates that at least one mutation was found in this gene/region in the community lysate sequence data. *PHROG based on phold annotations

Table S8 Loci with mutations in S-RIP4a isolates or communities coevolved or evolved with *Synechococcus* WH7803

| S-RIP4a locus_tag | Product | ISL [a] | T7-like Core | PHROG | PHROG Annotation | Functional Category | Coevolved WH7803 | | Evolved WH7803 | | | |
|---|---|---|---|---|---|---|---|---|---|---|---|---|
| | | | | | | | F [b] | G | 1 | 1 | 2 | 2 |
| | | | | | | | 8 | 7 | 1 | C | 1 | C |
| R4a070909_01 | hypothetical protein | ISL1 | | none | | | 1 | | | | | |
| IGR 5' R4a070909_08 | hypothetical protein | ISL1 | | none | | | 1 | | | | | |
| R4a070909_16 | DNA primase/helicase | | gp4 | 239 | DNA primase/helicase | RNA and nucleotide metabolism | | 1 | | | | |
| R4a070909_23 | ribonucleotide reductase | | | 237 | ribonucleotide reductas | RNA and nucleotide metabolism | 3 | 1 | | | | |
| R4a070909_38 | tail protein | | gp12 | 457 | tail protein | tail protein | | 1 | | | | |
| R4a070909_42 | internal virion protein | | gp16 | 6,651 | internal virion protein | head and packaging | 8 | 7 | 1 | X | 1 | X |
| R4a070909_43 | tail fiber protein | | gp17 | 29,288 | hypothetical protein | | 8 | 7 | 1 | X | 1 | X |
| R4a070909_46 | hypothetical protein | ISL3 | | none | | | 1 | | | | | |
| R4a070909_47 | hypothetical protein | ISL3 | | 5,939 | tail | tail protein | | 2 | | | | |
| R4a070909_49 | hypothetical protein | ISL3 | | 7802 | hypothetical protein | | | 1 | | | | |
| R4a070909_52 | hypothetical protein | ISL3 | | 12,220 | hypothetical protein | | 1 | 2 | 1 | X | 1 | X |
| R4a070909_56 | hypothetical protein | ISL3 | | none | | | | | 1 | X | 1 | X |
| R4a070909_59 | hypothetical protein | | | none | | | | | | | 1 | X |

[a] Genomic Island. [b] Letters or numbers in top row are the replicate designation. The second row is the number of isolates sequenced in each replicate, 'C' indicates that the community lysate was sequenced. The number in the columns represents the number of isolates within the replicate that possessed at least one mutation in the gene/region, an "X" in the column indicates that at least one mutation was found in this gene/region in the community lysate sequence data. *PHROG based on phold annotations

Table S9 Loci with mutations in S-RIP4b isolates or communities coevolved or evolved with *Synechococcus* WH7803

| S-RIM4b locus_tag | Product | ISL [a] | T7-like Core | PHROG | PHROG Annotation | Functional Category | Coevolved WH7803 | | Evolved WH7803 | | | |
|---|---|---|---|---|---|---|---|---|---|---|---|---|
| | | | | | | | F [b] | G | 1 | 1 | 2 | 2 |
| | | | | | | | 7 | 8 | 1 | C | 1 | C |
| IGR 5' R4b200813_01 | 5' hypothetical protein | ISL1 | | | | | 1 | | | X | | X |
| R4b200813_02 | hypothetical protein | ISL1 | | none | | | | 1 | | | | |
| R4b200813_17 | DNA polymerase | | gp5 | 15,971 | DNA polymerase | RNA and nucleotide metabolism | | 1 | | | | |
| R4b200813_19 | hypothetical protein | | | 27,006 | nblA | | 1 | 4 | | | | |
| R4b200813_20 | hypothetical protein | | | none | | | | 4 | | | | |
| R4b200813_21 | nucleotide pyrophosphohydrolase | | | 671 | nucleotide pyrophosphohydrolase | | | 1 | | | | |
| IGR 5' R4b200813_31 | 5' hypothetical protein | | | | | | 1 | | | | | |
| R4b200813_35 | tail protein | | gp11 | 446 | tail | tail protein | | 1 | | | | |
| R4b200813_36 | tail protein | | gp12 | 457 | tail | tail protein | 1 | 2 | | | | |
| R4b200813_39 | hypothetical protein | | gp15 | none | | | 1 | | | | | |
| R4b200813_41 | tail fiber protein | | gp17 | 29,288 | hypothetical protein | | 7 | 8 | 1 | X | 1 | X |
| R4b200813_46, 47, 48 | hypothetical proteins | ISL3 | | 17,104 | hypothetical protein | | 3 | 1 | | | | |
| R4b200813_48 | hypothetical protein | ISL3 | | 17,104 | hypothetical protein | | 2 | | | | | |
| R4b200813_51 | hypothetical protein | ISL3 | | none | | | 1 | | | | | |
| IGR 5' R4b200813_58 | 5' terminase large subunit | ISL3 | | | | | | 1 | | | | |
| R4b200813_58 | terminase large subunit | | gp19 | 38 | terminase large subunit | head and packaging | | 1 | | | | |
| R4b200813_59 | hypothetical protein | | | none | | | | | | X | | |
| IGR 5' R4b200813_60 | thymidylate synthase | | | 5' 85 | thymidylate synthase | RNA and nucleotide metabolism | 2 | | | | | |

[a] Genomic Island. [b] Letters or numbers in top row are the replicate designation. The second row is the number of isolates sequenced in each replicate, 'C' indicates that the community lysate was sequenced. The number in the columns represents the number of isolates within the replicate that possessed at least one mutation in the gene/region, an "X" in the column indicates that at least one mutation was found in this gene/region in the community lysate sequence data. *PHROG based on phold annotations
